# Dual color mesoscopic imaging reveals spatiotemporally heterogeneous coordination of cholinergic and neocortical activity

**DOI:** 10.1101/2020.12.09.418632

**Authors:** Sweyta Lohani, Andrew H. Moberly, Hadas Benisty, Boris Landa, Miao Jing, Yulong Li, Michael J. Higley, Jessica A. Cardin

## Abstract

Variation in an animal’s behavioral state is linked to fluctuations in brain activity and cognitive ability. In the neocortex, state-dependent control of circuit dynamics may reflect neuromodulatory influences including acetylcholine (ACh). While early literature suggested ACh exerts broad, homogeneous control over cortical function, recent evidence indicates potential anatomical and functional segregation of cholinergic signaling. Additionally, it is unclear whether states as defined by different behavioral markers reflect heterogeneous cholinergic and cortical network activity. We performed simultaneous, dual-color mesoscopic imaging of both ACh and calcium across the neocortex of awake mice to investigate their relationships with behavioral variables. We find that increasing arousal, categorized by different motor behaviors, is associated with spatiotemporally dynamic patterns of cholinergic release and enhanced large-scale network correlations. Overall, our findings demonstrate that ACh provides a highly dynamic and spatially heterogeneous signal that links fluctuations in behavior to functional reorganization of cortical networks.

## Introduction

Animals cycle through multiple waking brain states that profoundly influence patterns of neuronal activity, perception, and behavior^1-4^. Waking states can be categorized by a variety of cognitive and motor variables, including pupil dilation, facial movement, locomotion, arousal, and attention ^1, 5^. However, it is unclear whether variation in these parameters reflects different underlying brain network dynamics. A growing body of research suggests that arousal, measured using a variety of metrics including elevated motor activity, is associated with distinct alterations in local circuit operations within the neocortex, including changes in mean firing rates and decorrelation of the spike output of neighboring cells^3, 6-9^. Moreover, widefield, mesoscopic imaging^10^ has revealed broad representation of motor signals across the cortex^8^, with changes in behavioral state also linked to reorganization of functionally connected cortical networks^11, 12^. While desynchronization of local cortical network activity is a hallmark of transitions from periods of quiescence to periods of high arousal or motor output, it is unclear whether large-scale circuits spanning multiple cortical areas also exhibit changes in coordination across state transitions.

Classical views suggest that variation in neural activity associated with behavioral state fluctuations reflects the brain-wide, homogeneous influence of ascending neuromodulatory systems^13^. For example, cholinergic neurons in the basal forebrain send widespread projections throughout the neocortex that are thought to contribute to the effects of arousal, attention, and emotional valence on cortical dynamics^5, 14-19^. Application of ACh evokes desynchronization of local field potentials, de-correlates neuronal spiking, and enhances response gain, phenomena linked to enhanced attention and information representation^6, 19-28^. In addition, cholinergic neurons fire strongly to positive and negative reinforcement^29, 30^, resulting in reinforcement-related plasticity in the cortex^15, 31^. These studies suggest that highly salient environmental stimuli may produce uniform ACh signaling across the cortex to enhance global information processing and plasticity. However, recent electrophysiological and anatomical studies indicate substantial diversity in the firing patterns and axonal arborizations of individual cholinergic neurons, suggesting their output may instead be spatially and temporally heterogeneous in different cortical areas^7, 32-35^. Furthermore, the spatiotemporal relationships between cholinergic signaling and cortical neuronal activity across distinct behavioral states are unknown.

Monitoring cholinergic activity *in vivo* has typically required invasive probes with limited spatiotemporal resolution^36^. However, the development of receptor-based fluorescent indicators that directly report ACh binding has opened new avenues for the exploration of neuromodulation. Here, we used two-color mesoscopic imaging^10^ of the red-fluorescent calcium indicator jRCaMP1b^37^ and the green-fluorescent ACh indicator ACh3.0^38^ across the entire dorsal neocortex of the awake mouse to quantify the relationships between behavioral state, cortical activity, and cholinergic signaling. Our results demonstrate that, in contrast to traditional models, different behavioral states categorized by motor behavior are associated with distinct spatiotemporal patterns of ACh fluctuations and increased large-scale network synchronization. Overall, these findings re-shape our view of neuromodulation from a global state variable to a highly dynamic and spatially heterogeneous signal that links fluctuations in behavior to cortical network dynamics.

## Results

### Dual-color mesoscopic ACh and calcium imaging

To simultaneously monitor neuronal activity and cholinergic signaling in the neocortex of awake mice, we expressed the red fluorescent calcium indicator jRCaMP1b^37^ and the green fluorescent ACh sensor ACh3.0^38^ throughout the brain via neonatal injection of AAV vectors into the transverse sinus^11, 39^. This approach resulted in cortex-wide, uniform expression of both reporters (Figure S1), and we confirmed both *ex vivo* and *in vivo* that ACh3.0 specifically reported cholinergic signaling (Figure S2). We then performed mesoscopic imaging^10^ of both reporters through the intact skull of mice that were head-fixed and freely running on a wheel (Figure 1a, see Methods). Imaging was performed by strobing 575nm (jRCaMP1b), 470nm (ACh3.0), and 395nm (control) excitation light with an overall frame rate of 10 Hz per channel. ACh3.0 and jRCaMP1b images were co-registered, aligned to the Allen Common Coordinate Framework (CCFv3, Figure S2)^40^, and normalized (ΔF/F). There was no observable cross-talk between the green (ACh3.0) and red (jRCaMP1b) channels (Figure S2). We removed hemodynamic and motion-related artifacts using a novel regression approach that leverages spatial correlations in signal and noise and takes advantage of the reduced ACh-sensitive fluorescence of ACh3.0 when excited at 395nm^38^ (Figure S3, see Methods). This method outperformed conventional pixel-wise regression in its ability to remove stimulus-induced and spontaneous behavior-associated negative transients in green fluorescence corresponding to hemodynamic absorption^41, 42^ for both ACh3.0 and GFP (Figure S3).

**Figure 1.**
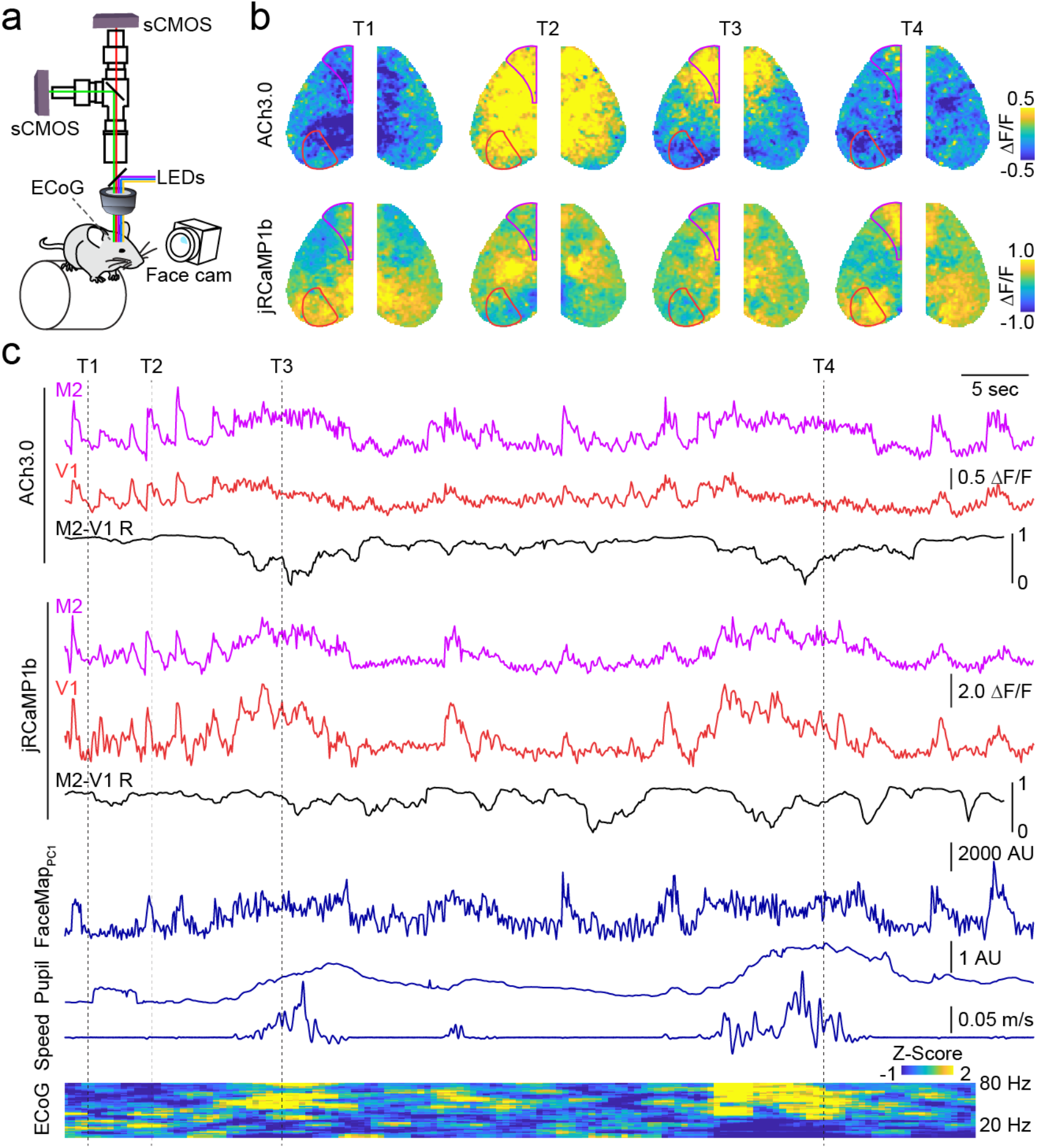
Spatiotemporal dynamics of cholinergic and neural activity in the neocortex. **a**, Schematic of the dual wavelength, widefield imaging setup. **b**, Example representative image frames showing fluctuations in cholinergic (ACh3.0, top row) and neural (jRCaMP1b, bottom row) activity across the dorsal cortical surface. Overlaid lines indicate Allen Brain CCFv3-derived parcellation of primary visual cortex (V1) and secondary motor cortex (M2). **c**, Time series for ACh3.0 and jRCaMP1b signals in V1 (red) and M2 (purple) parcels. Instantaneous (3 s window) Pearson’s correlation (R) between V1 and M2 is shown beneath in black. Simultaneous facial motion (FaceMap PC1), pupil area, running speed, and ECoG power spectrogram are shown below. Dashed lines indicate the times of image frames in (b).

Both ACh and calcium signals exhibited spatially heterogeneous, spontaneous fluctuations across the cortex (Figure 1b,c), illustrated by example data from two spatially distant cortical areas, primary visual (V1) and supplemental motor (M2) cortex. To investigate the relationship of this activity to behavior, we tracked the animal’s state using a combination of variables thought to reflect, in different ways, the animal’s overall level of arousal (Figure S4; Supplemental Table 1). Locomotion (wheel running speed) and pupil diameter have been repeatedly linked to variability in behavioral performance^4, 43-45^ and are associated with distinct patterns of local circuit dynamics^1, 3, 8, 46^. In addition, facial movements like whisking, which reflects the animal’s active exploration of its environment, are also closely coupled to cortical function^7, 9, 47^ and have been captured using FaceMap software to dimensionally reduce facial video data^9^. To complement imaging and behavioral data, we also recorded local circuit dynamics via electrocorticogram (ECoG) in a subset of experiments.

Overall, large fluorescence signal fluctuations occurred across cortical regions for both cholinergic and calcium signals and co-varied with changes in pupil dilation, locomotion, facial movement, and high-frequency ECoG activity (Figure 1c). Moreover, inter-areal correlations for both ACh and calcium were dynamic and tracked variation in behavioral variables. Spatially varied activity patterns in both channels were evident during specific periods of arousal (Figure 1, highlighted time frames), supporting a model in which ACh release is both dynamic and heterogeneous across cortical regions.

### Calcium and cholinergic signals encode behavioral state

To quantify the relationships between behavioral variables and fluorescence activity, we used a cross-validated linear regression model that included facial movements (FaceMap principal components 1-25, see Methods), locomotion speed, or pupil size as predictors of ACh or calcium fluctuations for each CCFv3-defined parcel. For ACh3.0, prediction accuracy (R^2^) differed significantly across cortical areas (p<0.001) and for different behavioral states (p<0.001, two way repeated measures ANOVA, n=6 mice per group), with facial movement providing the strongest prediction followed by locomotion and pupil variation (Figure 2a,c,e; face vs. locomotion: p = 0.003, face vs. pupil: p =0.001, locomotion vs. pupil: p = 0.065, post-hoc Least Significant Difference (LSD) tests). For jRCaMP1b, accuracy also differed significantly by area (p<0.001) and by state (p<0.001, two way repeated measures ANOVA, n=6 mice per group) and exhibited relative prediction for facial movement, locomotion, and pupil variation similar to Ach3.0 (Figure 2a,c,e; face vs. locomotion: p <0.001; face vs. pupil: p < 0.001; locomotion vs. pupil: p =0.068, post-hoc LSD tests). Using a Spearman’s rank correlation analysis, we also observed significant anterior-posterior gradients in the coupling between behavioral state and fluorescence activity that were inverted for cholinergic versus calcium signals (Figure 2b,d,f, see Methods), as ACh release and calcium signaling were best linked to behavioral state variables in frontal motor and posterior sensory areas, respectively. These results indicate that cholinergic release, like intracortical dynamics^8, 9^, encodes some forms of motor behavior and that distinct behavioral states are differentially coupled to neural activity. Statistical results for these and all subsequent analyses are listed in Supplemental Tables 2 and 3.

**Figure 2.**
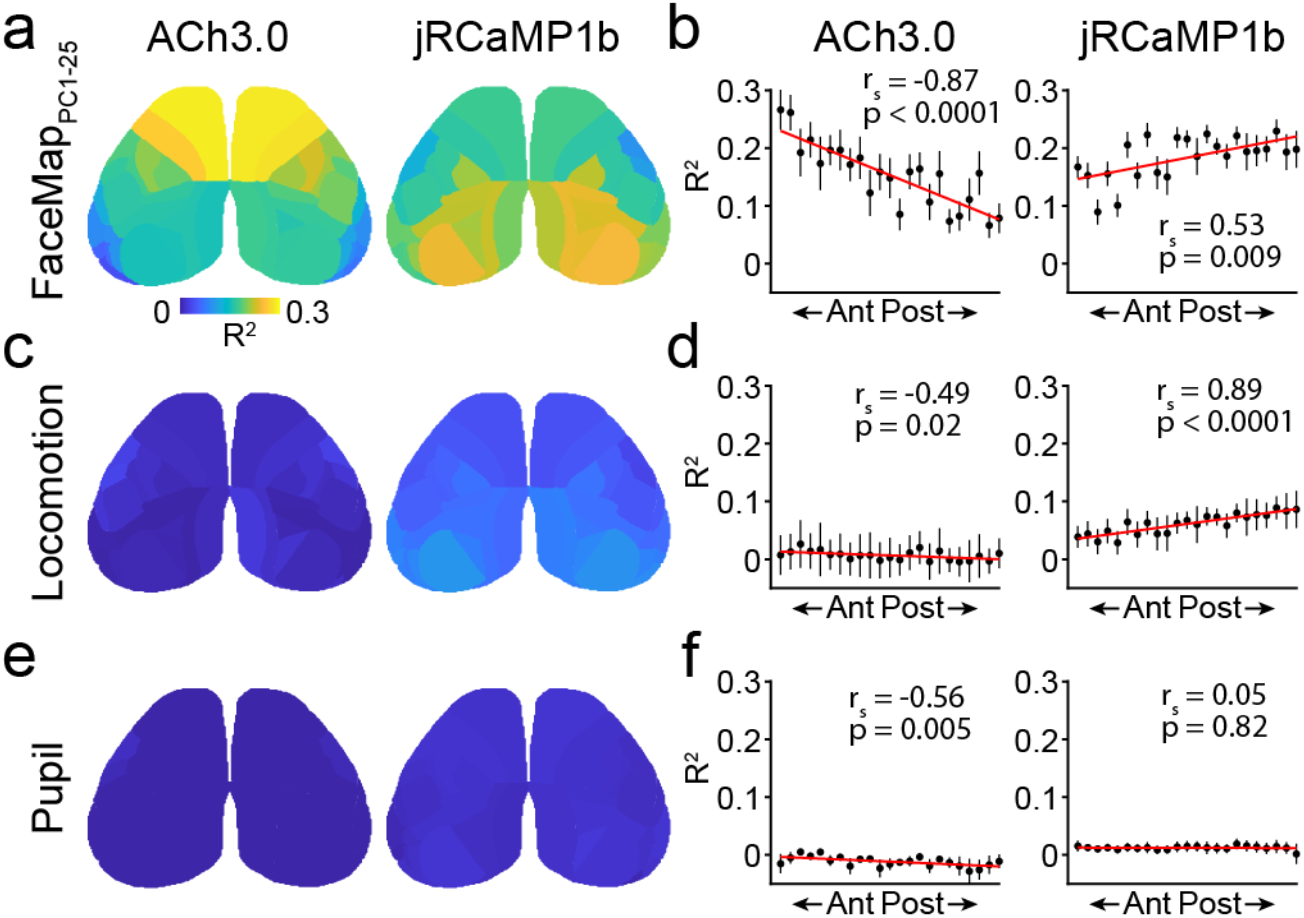
Differential coupling of behavioral variables to cholinergic and neural activity across the neocortex. **a**, Average spatial map (n=6 mice) showing ten-fold cross-validated explained variance (R^2^) of ACh3.0 (left) and jRCaMP1b (right) signals based on facial movements (FaceMap PC1-25). **b**, Allen Brain CCFv3 parcels from one hemisphere were ordered from anterior to posterior based on their center of mass. Cross-validated R^2^ values from each parcel (mean +/- SEM) are plotted against the parcel’s anterior-posterior position. r_s_ indicates Spearman’s rank order correlation coefficient for correlation between mean R^2^ and anterior-posterior rank across parcels. Line indicates linear fit for visualization. (**c, d**) and (**e, f**), same as in (a, b) for explained variance of ACh3.0 and jRCaMP1b signals based on locomotion and pupil area, respectively.

Given that facial movements and locomotion were the strongest predictors of both ACh and calcium signaling, we further explored these two motor-based indicators of arousal. State-dependent variation in brain activity might be accompanied by either transient or sustained changes in cholinergic and calcium signals that reflect underlying circuit dynamics. Therefore, we first identified timepoints corresponding to the onset of either locomotion or facial movements (Figure 3a, see Methods). Facial movement onset (quantified using FaceMap PC1) in the absence of locomotion was associated with desynchronization of local neuronal firing, as estimated by ECoG, and significant whole cortex-averaged transients in both ACh (0.64±0.09 ΔF/F, p=0.001, Student’s t-test, n=6 mice) and calcium (0.69±0.15 ΔF/F, p=0.006, Student’s t-test, n = 6 mice) signals (Figure S5a,b). Locomotion onset co-occurred with an increase in facial movement and local desynchronization and was also associated with transient increases in ACh (0.68±0.17 ΔF/F, p=0.010, Student’s t-test, n=6 mice) and calcium (1.50±0.11 ΔF/F, p<0.001, Student’s t-test, n=6 mice) signals (Figure S5a,b). ACh transients varied significantly across cortical areas (p<0.001) but were not significantly different between locomotion and facial motion onsets (p=0.734, two-way repeated measures ANOVA, n=6 mice per group). Calcium transients varied significantly across cortical areas (p<0.001) and states (p=0.005, two way repeated measures ANOVA, n=6 mice per group), with locomotion onset associated with the largest signal (Figure S5b). We also found a significant anterior-posterior gradient in the transient magnitudes for cholinergic but not calcium signals (Figure S5c).

**Figure 3.**
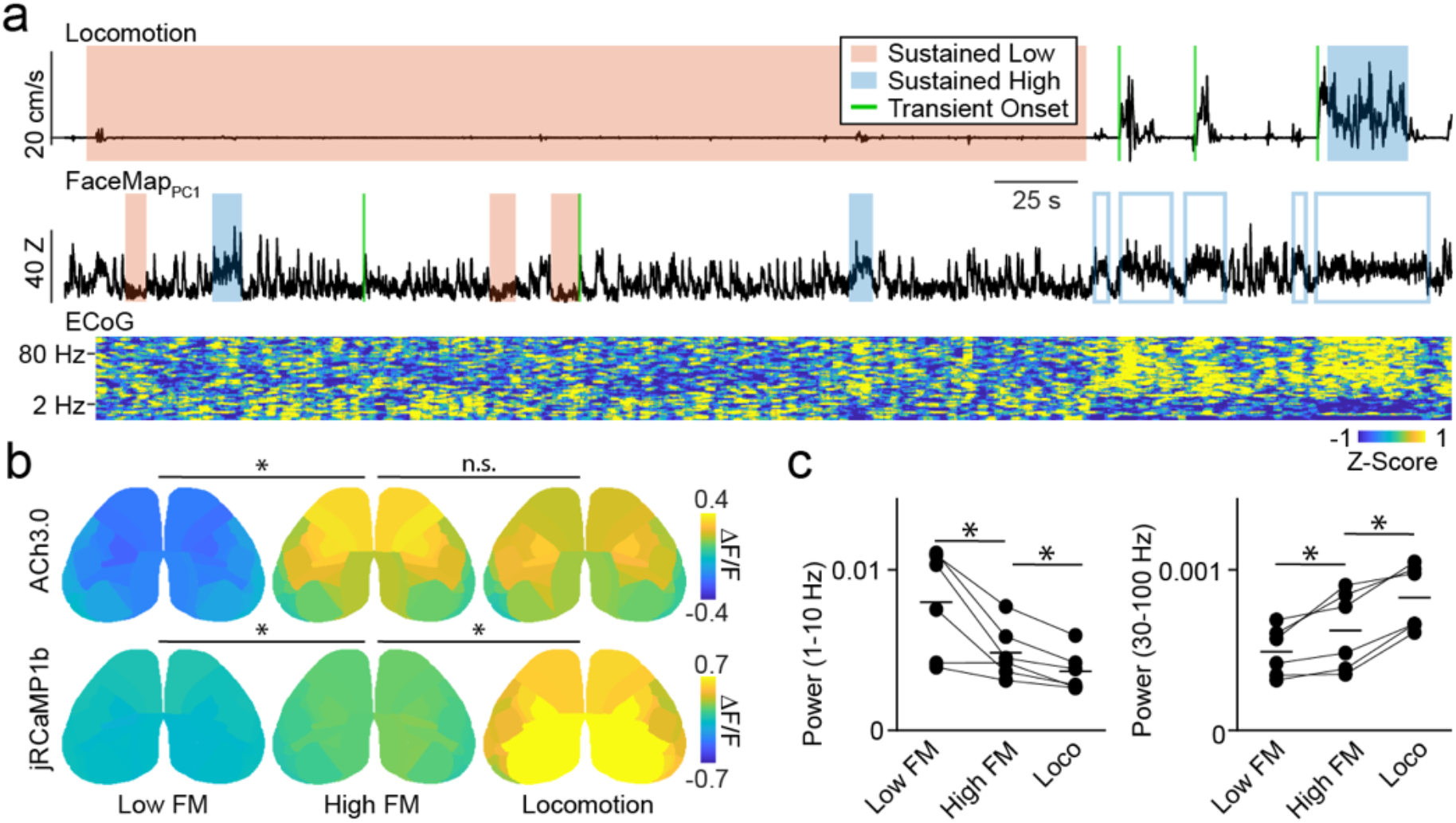
Cholinergic and neural signal heterogeneity during movement-defined behavioral states. **a**, Example locomotion (top) and FaceMap PC1 (bottom) traces from an experimental session. Periods of sustained low (red) or high (blue) motor activity are indicated by shaded regions and transitions from low-to high motor activity are delineated by green lines. Open blue boxes indicate high facial motion periods not included in analysis of sustained facial motor activity periods due to overlap with locomotion. **b**, Average spatial maps (n=6 mice) showing ACh3.0 (top) and jRCaMP1b (bottom) ΔF/F activity during sustained low facial motion, high facial motion, and locomotion states. *indicates p < 0.05 for post-hoc LSD comparisons following a significant (p < 0.05) main effect of behavioral state in repeated measures ANOVA. **c**, Individual animal values and population means for ECoG power in low (1-10 Hz) and high (30 – 100 Hz) frequency for three movement-defined sustained behavioral states. *indicates p < 0.05 for post-hoc LSD comparisons following a significant (p < 0.05) main effect of behavioral state in repeated measures ANOVA.

Next, we analyzed periods of sustained (>5 seconds) low or high motor activity (Figure 3a, see Methods). We observed that quiescence included epochs of both low and high facial movement whereas locomotion always co-occurred with high facial movement, suggesting that arousal might follow a progression from absence of motor activity, to facial motion, to locomotion. We examined the relationship of these three sustained states with cholinergic and calcium activity across cortical areas. ACh activity varied significantly across cortical areas (p<0.001) and differed by state (p=0.015, two way repeated measures ANOVA, n=6 mice per group). Sustained facial motion significantly elevated whole cortex Ach activity (Low facial motion: -0.13±0.04 ΔF/F, High facial motion: 0.18±0.08 ΔF/F; Low vs. High: p=0.046, post-hoc Least Significant Difference (LSD) test) but locomotion did not further increase the Ach signal (Locomotion: 0.16±0.06 ΔF/F; Locomotion vs High facial motion: p=0.717, post-hoc LSD test; Figure 3b). Similarly, sustained calcium activity varied significantly for cortical area (p<0.001) and state (p=0.005, two way repeated measures ANOVA, n=6 mice per group). Sustained facial movement and locomotion both increased whole cortex calcium activity (Low facial motion: -0.02±0.02 ΔF/F, High facial motion: 0.17±0.04 ΔF/F, Locomotion: 0.63±0.19 ΔF/F; Low vs. High facial motion: p=0.020, Locomotion vs High facial motion: p=0.044, post-hoc LSD test) (Figure 3b). As with transients, there was a significant anterior-posterior gradient in the differences across states measured with Spearman’s rank correlation for the cholinergic signal comparing sustained high and low facial movement and for calcium activity comparing sustained locomotion and high facial movement (Figure S5d). Sustained increases in both facial movement and locomotion were also associated with significant local circuit desynchronization, as evidenced by a decrease in low frequency power (main effect of state on 1-10 Hz: p = 0.001, repeated measures ANOVA; High vs. Low facial motion: p=0.022, Locomotion vs High facial motion: p=0.004, post-hoc LSD tests) and an increase in high frequency power (main effect of state on 30-100 Hz: p < 0.001, repeated measures ANOVA; High vs. Low facial motion: p=0.048, Locomotion vs. High facial motion: p=0.002, post hoc LSD tests) in the ECoG (Figure 3c). Notably, we observed a similar relationship between behavioral state and ACh signaling by carrying out mesoscale calcium imaging of GCaMP6s-expressing cholinergic fibers (Figure S6). Thus, these results demonstrate that facial movement is coupled to robust increases in ACh release that is greatest in frontal regions. In contrast, intracortical activity measured via calcium signals is most robustly increased by locomotion in posterior cortical regions.

### State-dependent modulation of cortical network architecture

Behavior is thought to rely on large-scale coordination of activity between cortical areas^48, 49^. Therefore, we next examined the spatiotemporal relationships of both ACh and calcium signals by measuring pairwise correlations between cortical regions. We focused on sustained periods of low facial movement, high facial movement, and locomotion, allowing us to test whether these behavioral states with different average increases in fluorescence signals also correspond to distinct changes in network organization. We found that increased facial movement was associated with significant increases in between-area pairwise correlations in anterior somatosensory and motor areas for ACh signal and broad, significant increases for calcium signal (Figure 4a,b, adjusted p<0.01, Benjamini-Yekutieli FDR corrected permutation test for each pair, n=6 mice). In marked contrast, locomotion was associated with a reduction in the between-area correlations of cholinergic activity that was significant across the cortex (Figure 4a, adjusted p<0.01, Benjamini-Yekutieli FDR corrected permutation test for each pair, n=6 mice). Locomotion also occurred with modest decreases in correlation for calcium signals that were significant primarily across posterior cortical areas (Figure 4b). We note that the limited number of pixels present for the most lateral CCFv3 parcels likely decreases the robustness of correlation measurements for these areas. Overall, our results suggest that moderate levels of arousal (i.e., facial movement without locomotion) generally enhance the coordination of ACh release in the cortex, but further increases in arousal (i.e., locomotion) profoundly decorrelate cholinergic signals, emphasizing the spatiotemporal independence of cholinergic modulation. Moreover, in contrast to the local network decorrelation associated with enhanced arousal (observed in the ECoG data), we find that facial movement is linked to significantly increased correlation across large-scale networks as measured by calcium imaging.

**Figure 4.**
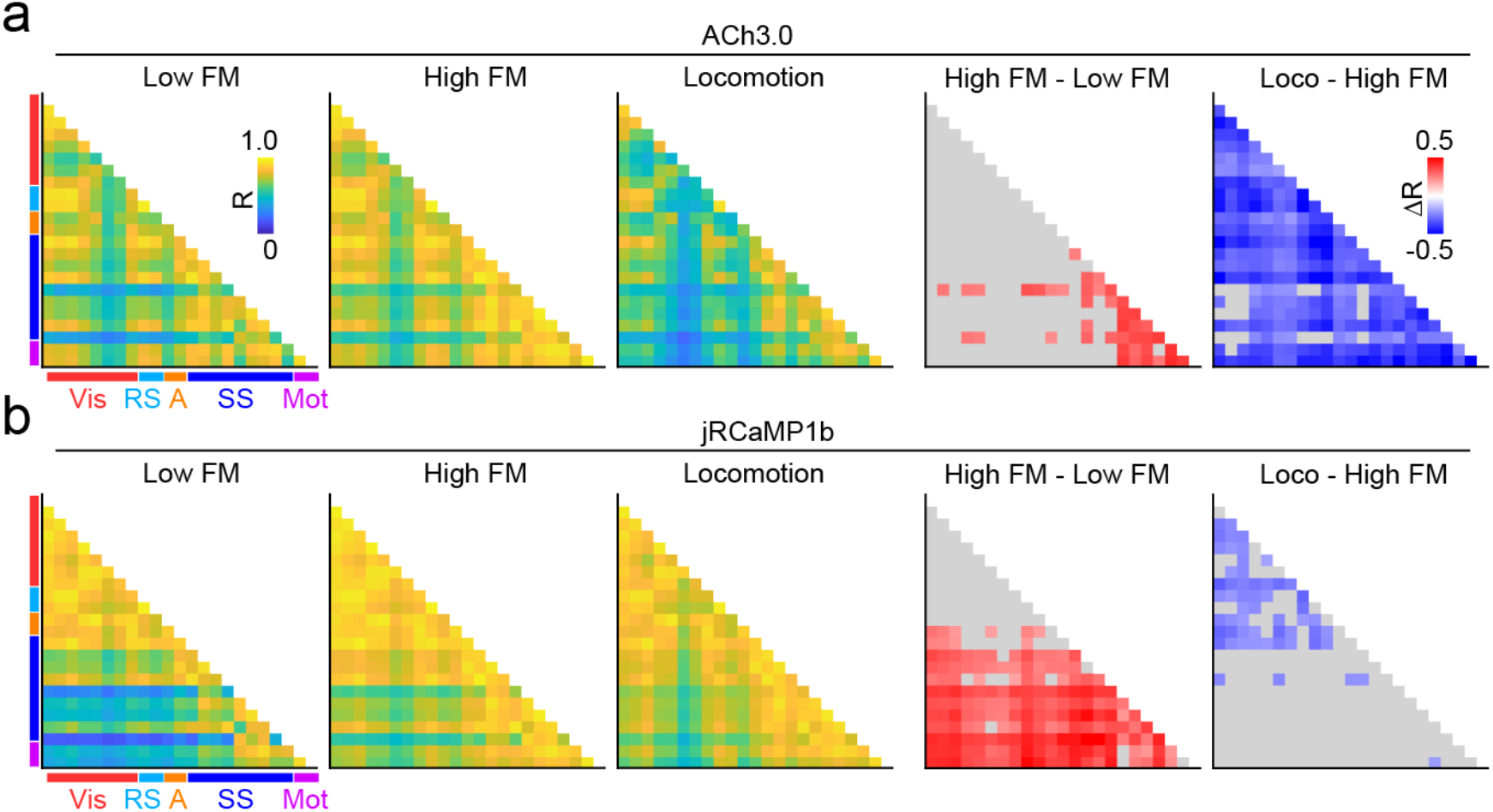
State-dependent variation in spatial correlations of cholinergic and neural activity. **a**, Average peak cross-correlation matrices (n=6 mice) for cholinergic activity during three movement defined behavioral states: low facial motion, high facial motion, and locomotion, measured pairwise for CCFv3-defined cortical parcels in the left hemisphere (1st, 2nd, and 3rd columns). Average differences in correlation between high and low facial motion states (4th column) and between locomotion and high facial motion states (5th column) are shown for significant pairs (adjusted p<0.01, Benjamini-Yekutieli FDR corrected permutation test). Non-significant pairs are shown in gray. Vis, Visual; RS, retrosplenial; A, Auditory; SS, somatosensory; Mot, Motor. **b**, As in (a) for calcium (jRCaMP1b) activity.

Increases in cholinergic signaling have been linked to enhanced local neuronal firing and response gain^6, 19, 25, 50, 51^. We therefore investigated whether ACh and calcium signals are coupled to each other within individual cortical areas and whether this relationship varies with behavioral state. Sustained facial movement was associated with a significant increase in correlation between the two signals broadly across the cortex (Figure 5a-b, adjusted p<0.01, Benjamini-Yekutieli FDR corrected permutation test for each pair, n=6 mice). In striking contrast, sustained locomotion was associated with a significant reduction in the correlation between the two channels that was restricted to the frontal cortex (Figure 5a-b, adjusted p<0.01, Benjamini-Yekutieli FDR corrected permutation test for each pair, n=6 mice). These data suggest that cholinergic coupling to neuronal activity varies with both cortical area and behavioral state. Overall, our findings demonstrate a spatially compartmentalized relationship between cholinergic modulation and cortical activity that is diversely dependent on transitions between distinct behavioral states.

**Figure 5.**
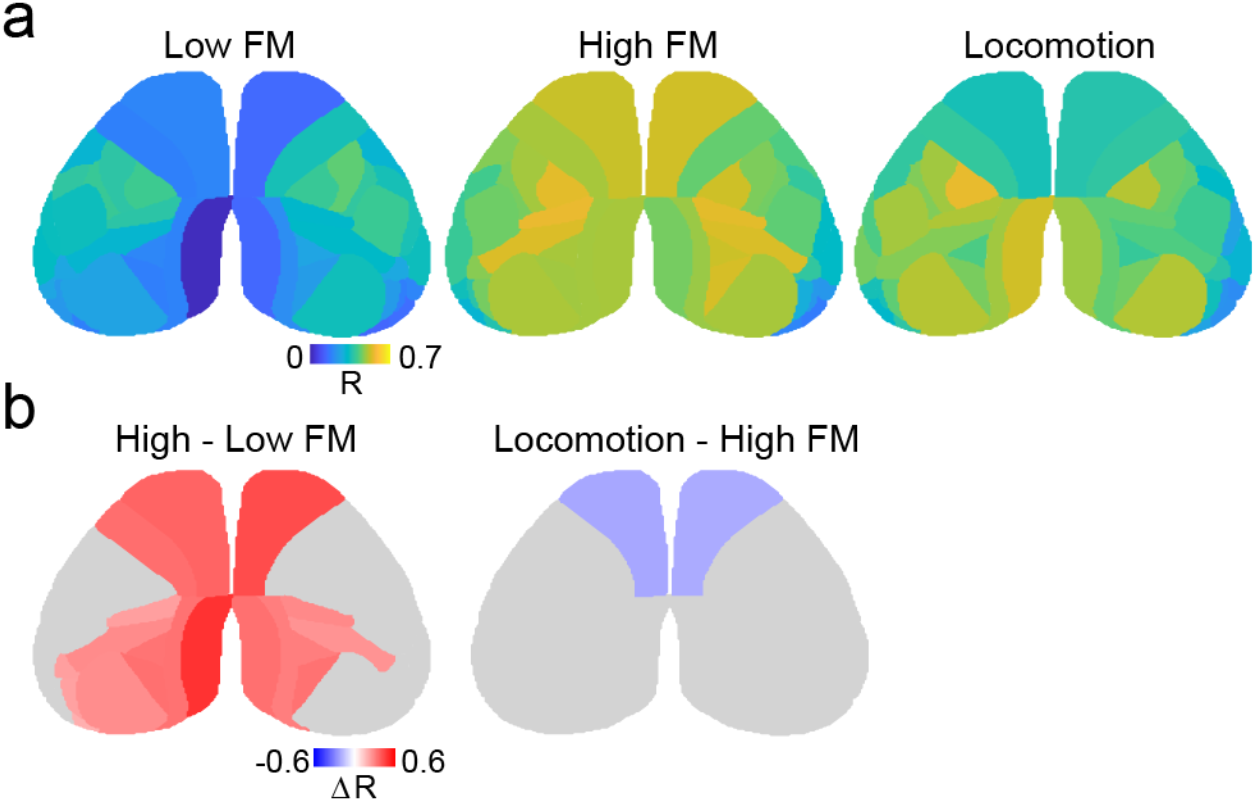
State-dependent spatial variation in correlations between cholinergic and calcium signals. **a**, Average spatial maps (n=6 mice) showing peak cross-correlation coefficients between ACh3.0 and jRCaMP1b activity for each cortical parcel during three movement defined behavioral states: low facial motion, high facial motion, and locomotion. **b**. Average difference in correlation between high and low facial motion states (left) and between locomotion and high facial motion states (right) is shown for significant parcels (adjusted p<0.01, Benjamini-Yekutieli FDR corrected permutation test). Non-significant parcels are shown in gray.

## Discussion

Despite a long history of investigation into the role of cholinergic modulation in the neocortex, experimental challenges to large-scale simultaneous monitoring of ACh release and neural activity *in vivo* have limited our understanding. Indeed, consistent observations that ACh is coupled to salient behavioral cues across sensory modalities might suggest this modulatory pathway serves as a global, homogeneous influence throughout the cortex^13, 52^. Here, we challenged this view using a combination of mesoscopic imaging and newly developed reporters of both calcium and ACh in behaving mice. Our results clearly demonstrate that cholinergic signaling exhibits substantial spatiotemporal fluctuations that are differentially related to distinct behavioral states, defined by patterns of motor activity.

Both facial movements and locomotion are associated with significant, broad increases in ACh release that are enhanced for anterior versus posterior areas. This anterior-posterior gradient could map onto slightly differential cholinergic projection densities in the frontal versus parietal and occipital areas, to rough antero-posterior and medio-lateral projection topography from the basal forebrain, and/or to different ratios of basal forebrain cholinergic to non-cholinergic (GABAergic) projections for frontal and posterior cortical areas^32, 53, 54^. Interestingly, we find that facial movement is coupled to significantly synchronized ACh release across the anterior cortex. In contrast, locomotion, which co-occurs with facial movement, is coupled to robust, cortex-wide desynchronization of cholinergic activity. This result emphasizes the fundamental independence of cholinergic signaling across different cortical areas and is consistent with anatomical and electrophysiological heterogeneity within populations of cholinergic projection neurons in the basal forebrain^32, 34, 35, 55, 56^. We speculate that information about behavioral states is relayed broadly to the entire basal forebrain, which then propagates independently via discrete channels to selective areas of the cortex. Such a system would allow independent, behavior-specific modulation of cortical subnetworks. We also note that ACh can be released locally by subpopulations of cortical interneurons, providing an additional mechanism for heterogeneous cholinergic modulation^57^.

Motor behavior is closely linked to fluctuations in the activity of neurons across the neocortex, suggesting that the representation of movement is a fundamental feature of cortical networks^8, 9, 58^. Here, we find similarly that facial movements and locomotion are represented in ACh signals. Regression analyses suggest that facial motor information is most prominent, contrasting somewhat with recent observations showing that cholinergic axonal activity is strongly coupled to pupil dynamics^46^. We note that, unlike axonal calcium imaging, ACh3.0 reports cholinergic release and the transfer function between presynaptic calcium and transmitter exocytosis *in vivo* is poorly characterized^59^. In this way, our approach is similar to electrochemical measurement of cholinergic release^60^ but capable of stable, longitudinal monitoring across the entire dorsal cortex in the absence of invasive probes. Importantly, none of these methods provide information regarding the physiological extent of ACh reception or cell type-specific downstream signaling, which ultimately requires tools for monitoring both postsynaptic biochemical activity and spike output at cellular resolution^61^.

In the neocortex, both motor activity and cholinergic signaling have been linked to the desynchronization of local circuits and the decorrelation of spiking between neighboring neurons^6, 22, 23, 25, 62^. Such electrophysiological changes are suggested to enhance the representation of information, particularly under demanding cognitive regimes such as focused attention^63^. However, the coordination of sensory, motor, and cognitive networks is also thought to be necessary for behavior^1, 8, 9, 64^. Here, we find the surprising result that increasing motor activity and ACh release are associated with significant increases in long-range, intra-cortical correlations even though local circuits are decorrelated.

The appreciation for behavioral state as a critical determinant of cortical function and behavior has only increased in recent years, although the diversity of studies has left a number of ambiguities unresolved. For example, variations in arousal, attention, locomotion, whisking, facial movements, and pupil diameter have all been linked to cortical dynamics and modulation of perceptual ability ^1-4, 9, 45^. The extent to which these categorizations reflect distinct or overlapping mechanisms and computations remains largely unknown. Indeed, previous work found that arousal without locomotion and arousal accompanied by locomotion exert distinct influences on local cortical activity patterns, but both enhance sensory encoding^3^. Nevertheless, our findings here underscore the hypothesis that waking comprises diverse behavioral states characterized by specific patterns of neuromodulatory dynamics^3, 13^. Indeed, our data suggest that arousal and neuromodulation may follow a progression that parallels motor activity, first manifesting with facial movement and increasing with locomotion (Figure 6). In this model, moderate arousal drives increased ACh release and cortical activity that is coupled to a decorrelation of local circuits and enhanced large-scale synchrony. Further arousal associated with locomotion drives desynchronized ACh release, local circuit decorrelation, and global correlation of broad intracortical networks. The causal links between these dynamics remain to be determined. However, the advent of multi-color fluorescent reporters, novel combinations of imaging modalities and electrophysiology^11, 12, 38, 65^, and further refinement of behavioral analyses^9, 66^ will open further avenues into resolving these issues.

**Figure 6.**
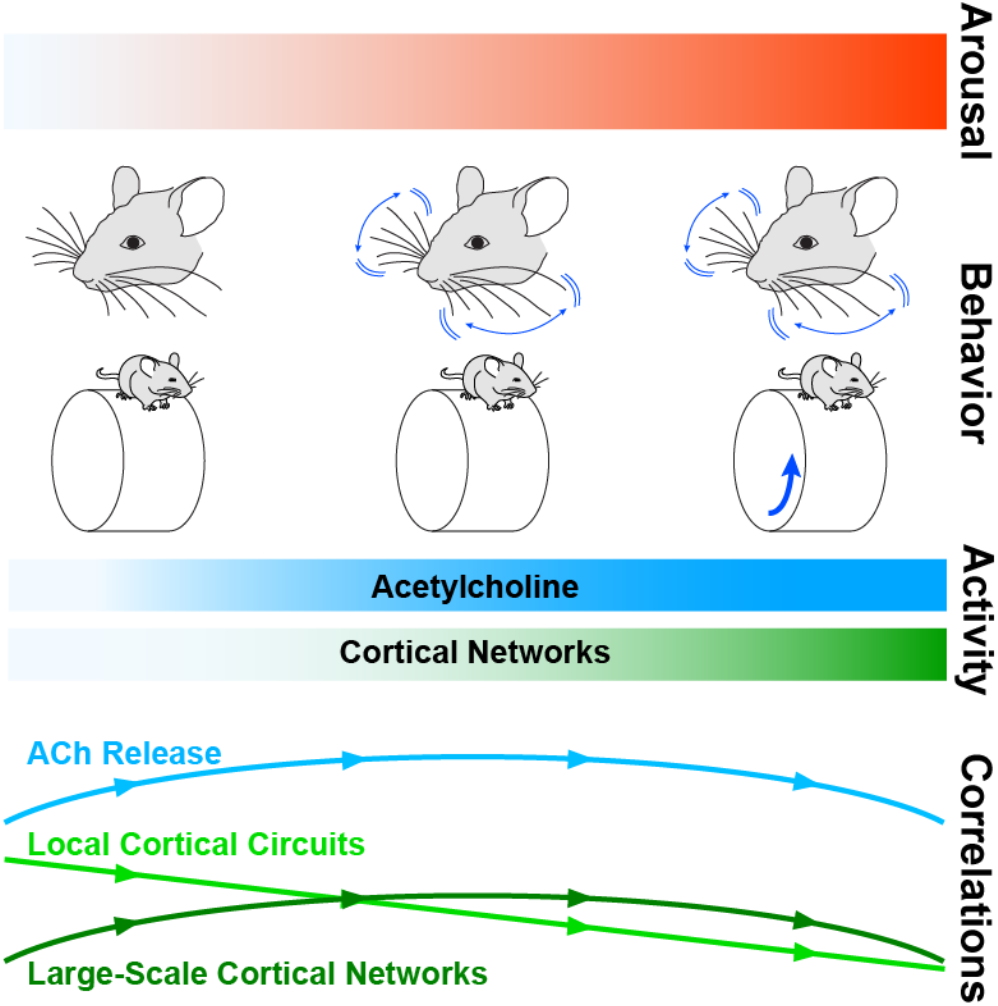
Relationship between arousal, cortical activity, and network synchrony. Arousal is associated with distinct patterns of motor activity, marked by 1) elevated facial motion energy arising from movements of whiskers and facial muscles and 2) locomotion, which always co-occurs with facial movement. Increasing arousal corresponds to elevated neuronal activity and ACh release across the neocortex. With increased arousal, correlations within local cortical circuits measured via ECoG progressively decrease. In contrast, correlations within large-scale networks and of Ach release across the cortex measured with mesoscopic calcium imaging increase during moderate arousal associated with high facial motion energy but decrease with further increases in arousal with locomotion.

In conclusion, we provide novel evidence for the idea that movement is broadly represented in the cortex, not only in local circuit activity^8, 9^ but also in the dynamics of cholinergic modulation. Furthermore, cholinergic output during movement is spatially structured and non-uniform in the cortex. Additional studies are required to determine whether these common patterns reflect movement-related ascending regulation of cortical networks or top-down, state-dependent control of basal forebrain output^13^. In either case, we propose the broad hypothesis that the fine-tuned orchestration of behavioral and cortical state dynamics requires coordinated coupling of neuronal activity and cholinergic modulation within distinct, spatially heterogeneous networks of the neocortex.

## Author Contributions

SL, AHM, MJ, YL, MJH, and JAC designed the experiments. SL and AHM collected the data. SL, AHM, HB, and BL analyzed the data. SL, AHM, MJH, and JAC wrote the manuscript.

## Acknowledgements

The authors thank all members of the Higley and Cardin laboratories for helpful input throughout all stages of this study. We thank Rima Pant for generation of AAV vectors. We thank Daniel Barson, Gal Mishne, and Ronald Coifman for helpful discussions regarding the analysis of the data. We thank Quentin Perrenoud for providing the locomotion changepoint analysis code. We thank the GENIE Project for jRCaMP1b plasmids. This work was supported by funding from the NIH (MH099045 and MH121841 to MJH, EY022951 to JAC, MH113852 to MJH and JAC, EY031133 to AHM, EY026878 to the Yale Vision Core), an award from the Kavli Institute of Neuroscience (to JAC and MJH), a Simons Foundation SFARI Research Grant (to JAC and MJH), a Swebilius Foundation award (to JAC and MJH), support from the Ludwig Foundation (to JAC), a BBRF Young Investigator Grant (to SL) and an award from the Swartz Foundation (to HB).

## Data Availability Statement

The full datasets generated and analyzed in this study are available from the corresponding authors on reasonable request. All values found in Figures 4 (correlations between pairs of parcels) and 5 (correlations between signals within a parcel) have been made available as an attached excel file.

## Code Availability Statement

Custom written MATLAB scripts used in this study are available from the corresponding authors on reasonable request.

## Supplemental Figures and Figure Legends

**Figure S1.**
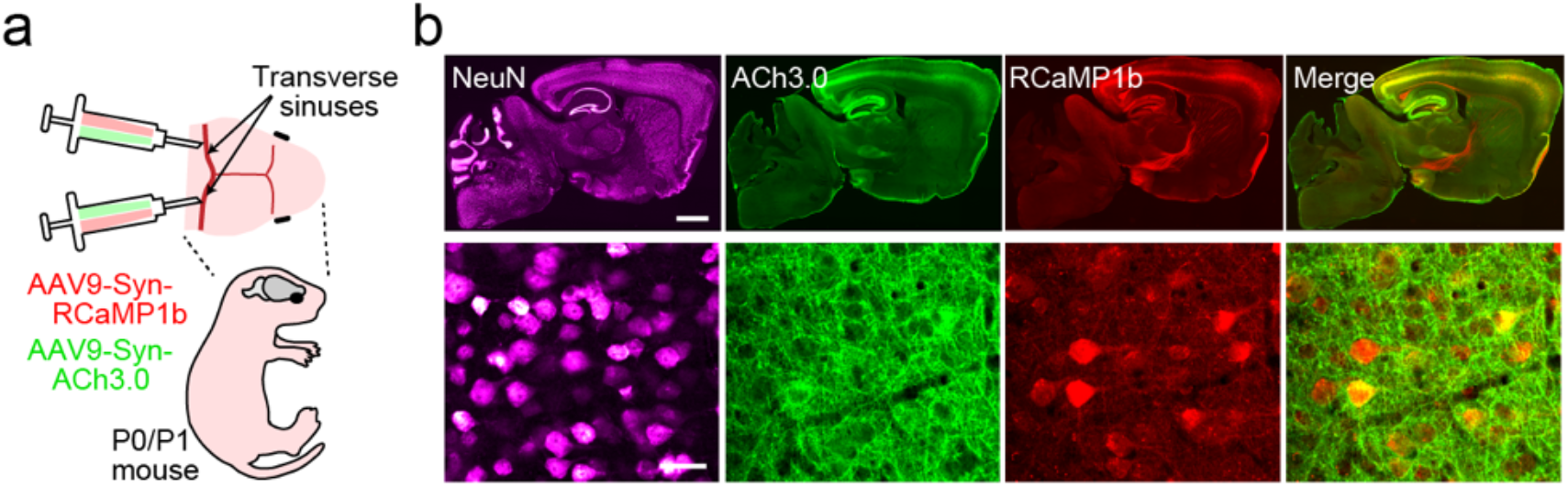
Viral expression of the cholinergic reporter ACh3.0. **a**, Schematic of the neonatal sinus injection approach. **b**. Example sagittal widefield (top) and confocal (bottom) fluorescent images from an adult mouse expressing jRCaMP1b (red) and ACh3.0 (green) following neonatal virus injection. Scale bars; 1mm top, 25 µm, bottom.

**Figure S2.**
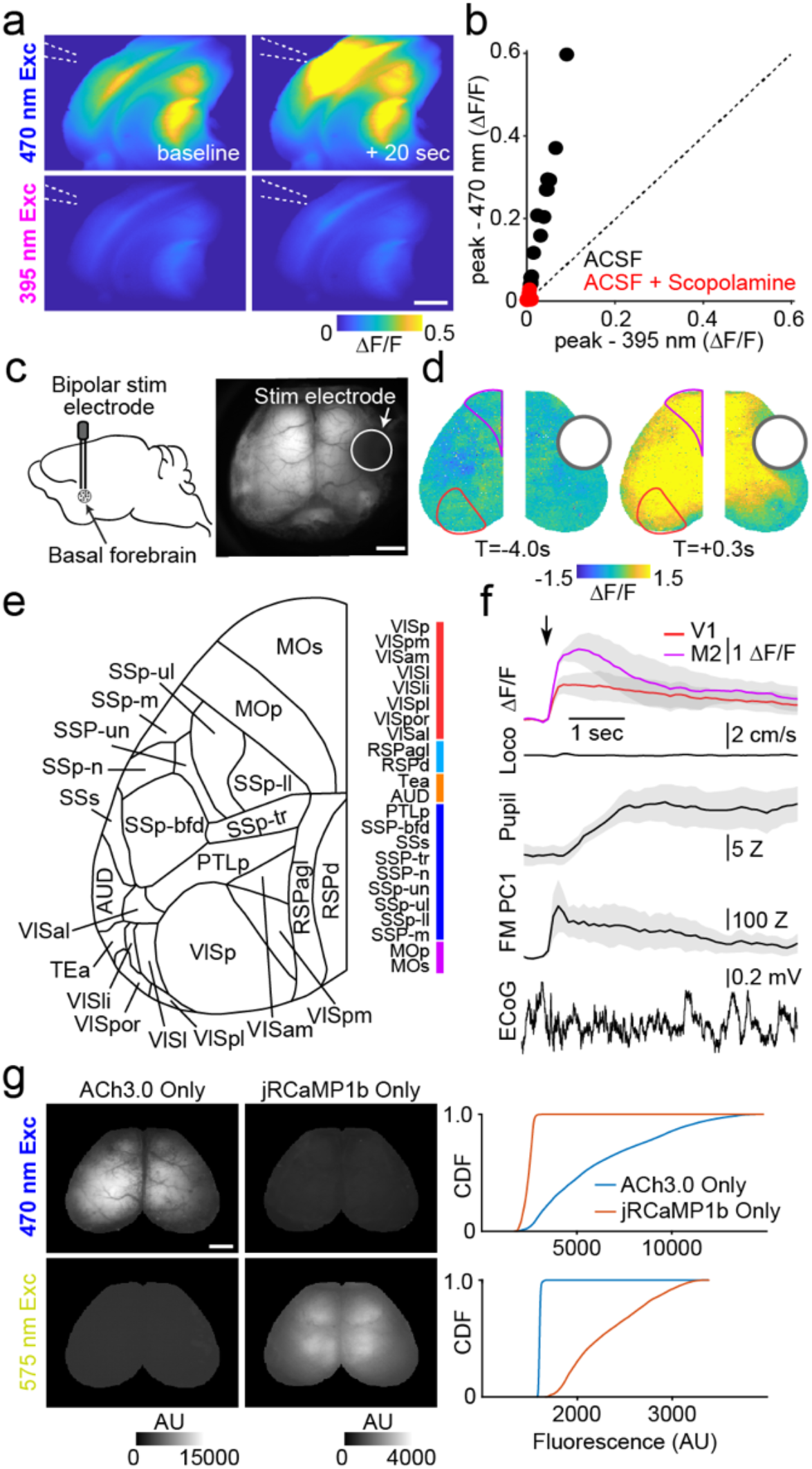
*ex vivo* and *in vivo* validation of mesoscopic ACh3.0 signals. **a**, Example brain slice from an ACh3.0-expressing mouse imaged with 470nm (blue) and 395 (violet) excitation light on interleaved frames. Images show fluorescence at baseline (left) and 20 second after carbachol (20 μM) was puffed onto the slice (right). Scale bar; 1mm. **b**, Peak carbachol-evoked ΔF/F responses for 395nm versus 470nm excitation, in control ACSF or in the presence of scopolamine (10 μM). **c**, Schematic showing bipolar stimulation electrodes implanted in the basal forebrain (left) and corresponding mesoscopic image showing baseline fluorescence (right). Scale bar; 1mm. **d**, Example pixel-wise spatial maps of ΔF/F ACh3.0 signal averaged across trials before (left) and after (right) electrical stimulation of the basal forebrain. **e**, Schematic illustrating Allen CCFv3 parcels, corresponding abbreviations, and color codes used in main figures. **f**, Mean +/- SEM (n=3 mice) ACh3.0 signal activity evoked by basal forebrain electrical stimulation for V1 (red) and M2 (purple) Allen CCFv3 parcels indicated in e. Corresponding Mean +/- SEM changes in locomotion, pupil, facial movement (FaceMap PC1), and an example ECoG trace from a single stimulation are illustrated below. **g**, Example images showing fluorescence under 470 nm excitation light (top) and 575 nm excitation light (bottom) in an ACh3.0-expressing mouse (left) and a jRCaMP1b-expressing mouse (right). Cumulative distribution plots of the pixel intensities from the example frames under both illumination conditions are shown on the far right.

**Figure S3.**
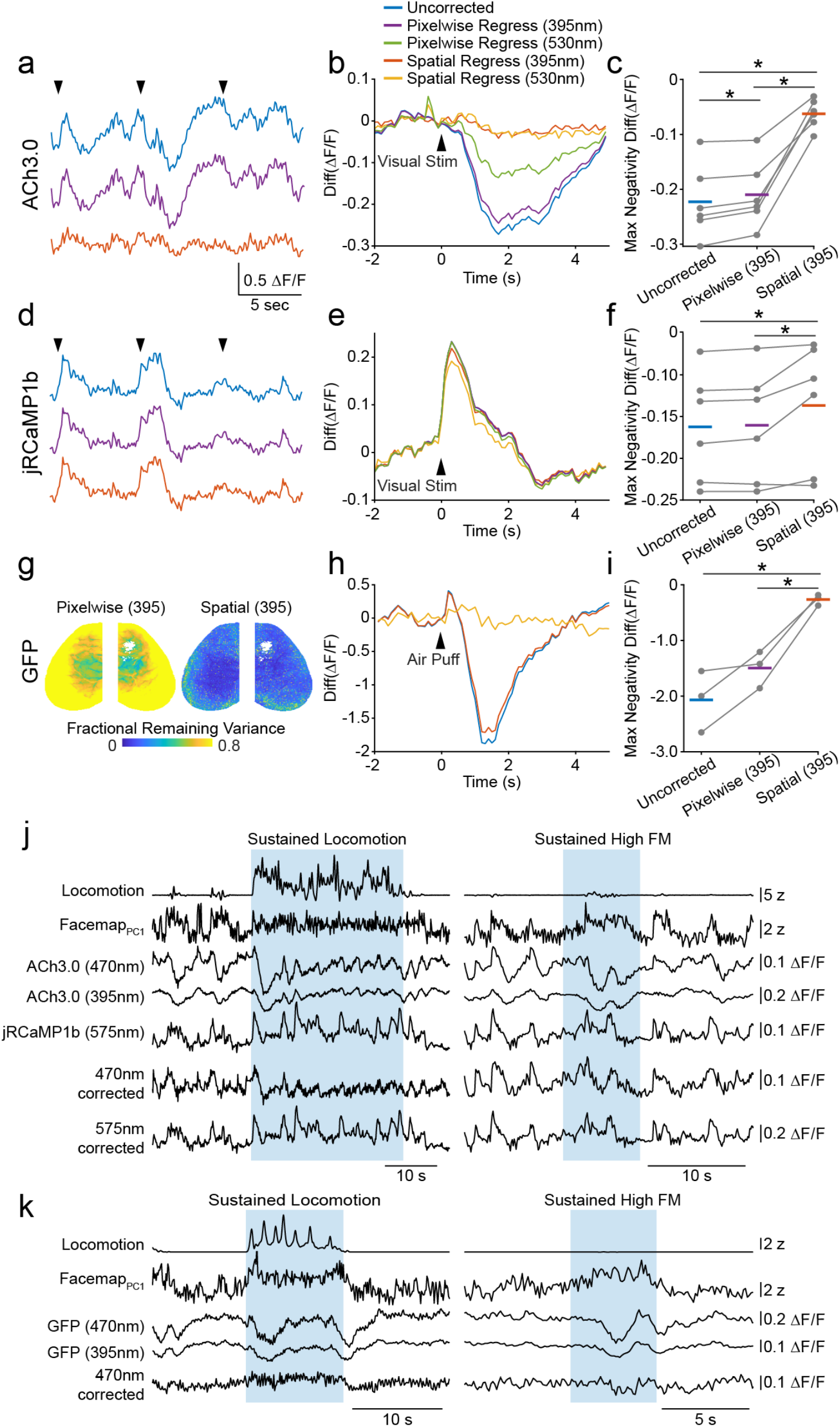
Spatial regression for correction of hemodynamic artifacts in mesoscopic data. **a**, ACh3.0 signals measured in V1 during presentation of drifting grating stimuli (arrows), illustrating different hemodynamic correction methods. **b**, Average ΔF/F ACh3.0 activity evoked by visual stimulation at times indicated in (a). Traces are for uncorrected fluorescence (blue) and images corrected using pixel-wise regression of 395nm fluorescence data (purple) or 530 nm back-scatter data (green), or spatial regression of 395 nm (red) or 530 nm (orange) data. **c**, Individual animal and population mean (n=6 mice) values for visually-evoked ΔF/F negativity (hemodynamic artifact) for the different regression methods. * indicates p<0.05, post hoc LSD tests following repeated measures ANOVA. **d-f**, as in (a-c) for jRCaMP1b data (n=6 mice) in V1. **g**, Pixel-wise variance remaining from imaging of a GFP-expressing mouse following hemodynamic correction with pixel-wise (left) or spatial (right) regression of 395nm fluorescence data. **h**, Average GFP fluorescence in V1 evoked by air-puff stimulus to the animal’s flank. Traces are for uncorrected fluorescence (blue) and images corrected using pixelwise (purple) or spatial (red) regression of 395nm data. **i**, Individual animal and population mean (n=3 mice) values for air-puff-evoked ΔF/F negativity for the different regression methods. * indicates p<0.05, post hoc LSD tests following repeated measures ANOVA. **j**, Example behavioral and neural data time series from a period of sustained locomotion (left) and sustained high facial motion activity (right) in a dual-expressing mouse. Neural data are V1 traces from ACh3.0 under 470 nm and 395 nm excitation light and jRCamp1b under 575 nm excitation light before and after hemodynamic correction using spatial regression of the 395 nm fluorescence. Correction substantially reduces large negative deflections especially in the ACh3.0 signal. **k**, Same as in (j) for an example GFP-expressing mouse.

**Figure S4.**
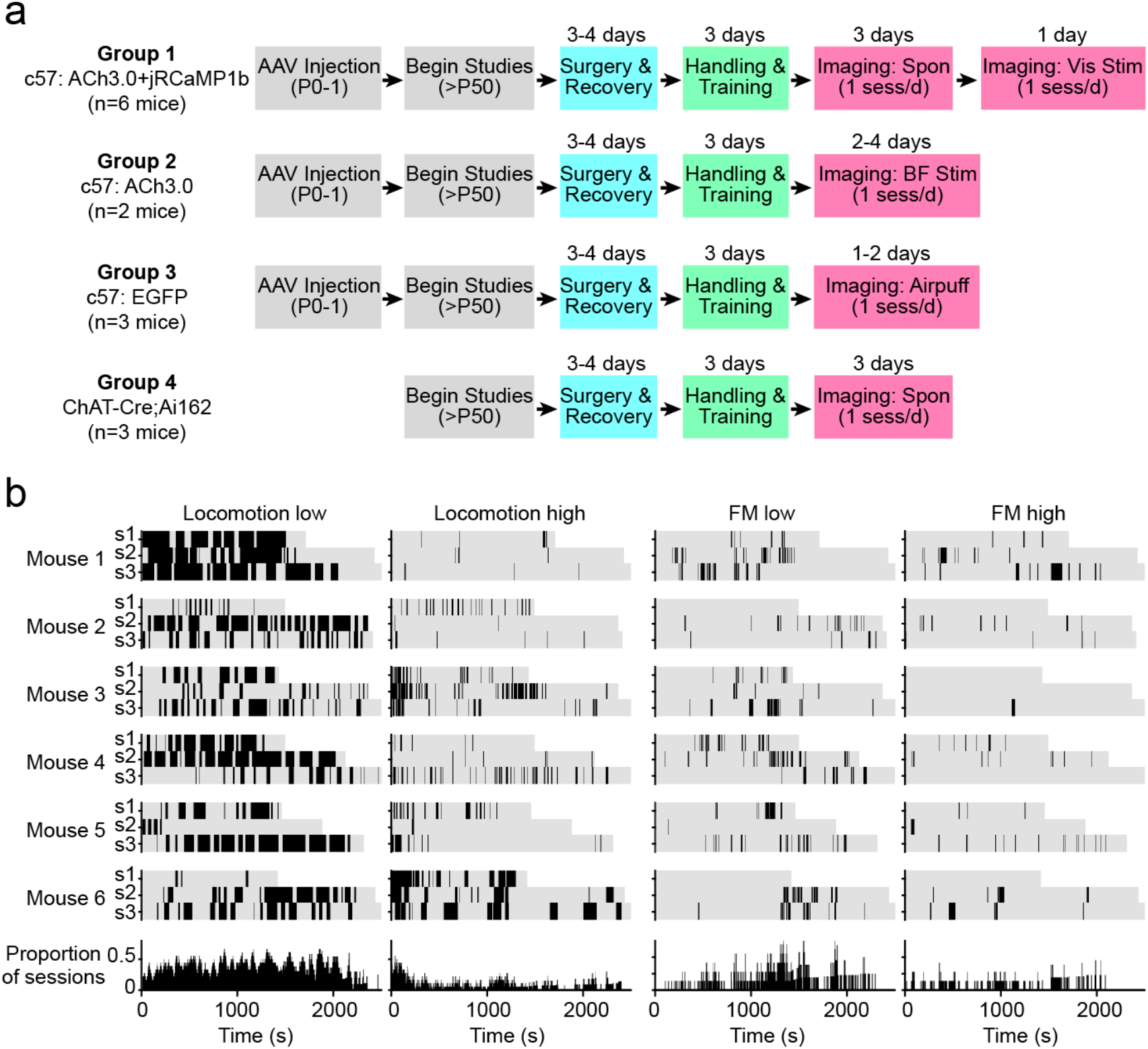
Experimental timeline and distribution of behavioral states. **a**, Experimental timeline and group schematic for all animals used in the study. **b**, Behavioral raster plots indicating periods of sustained low and high locomotion and sustained low and high facial motion activity across all imaging session in the six dual ACh3.0/jRCaMP1b mice comprising Group 1. Below are histograms indicating the distribution of the sustained states over time.

**Figure S5.**
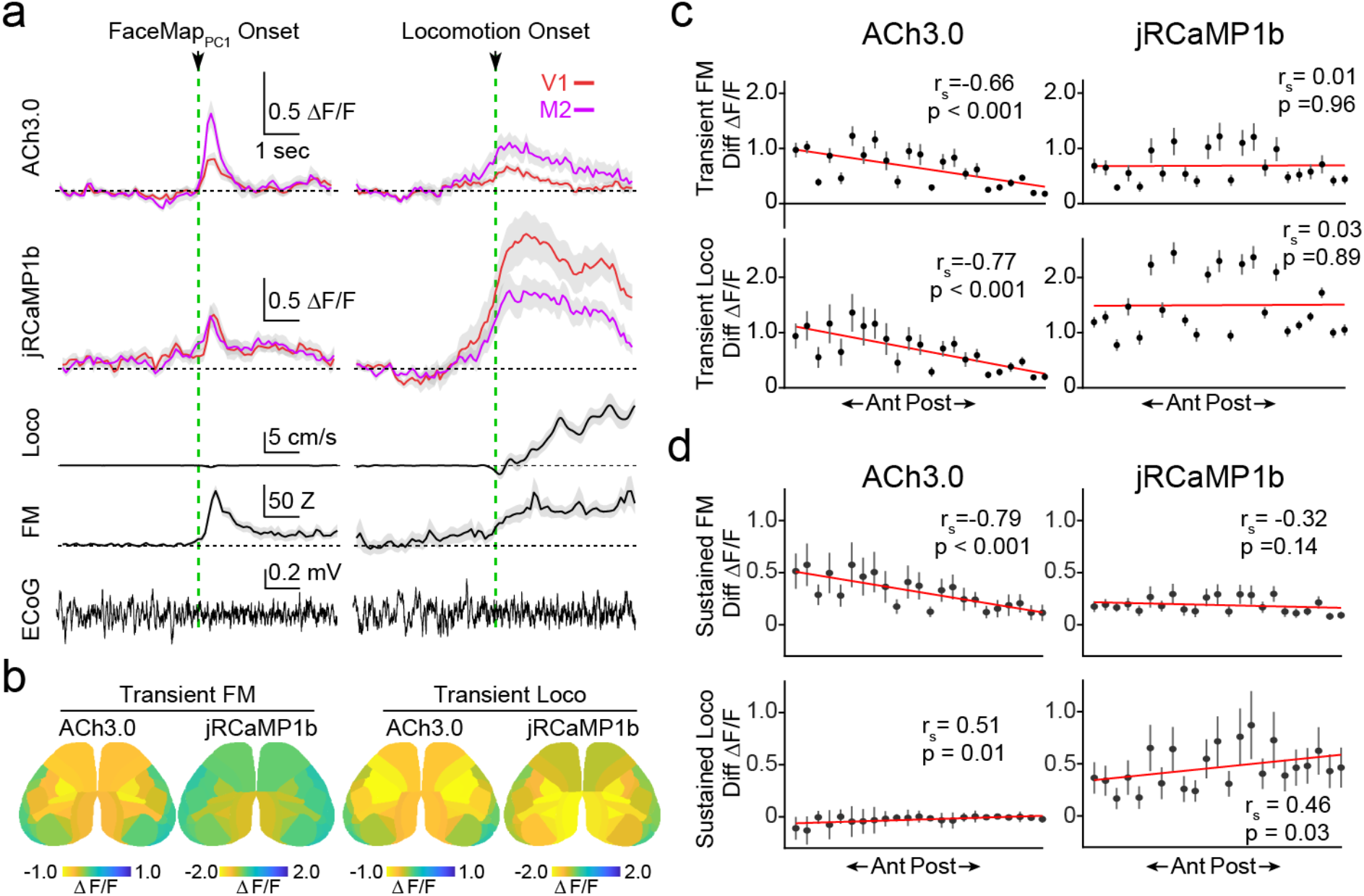
Spatial heterogeneity in cholinergic and calcium signaling during movement-defined transient and sustained behavioral states. **a**, Traces show trial-averaged activity for V1 (red) and M2 (purple) aligned to FaceMap PC1 (left) and locomotion (right) transition points for one session, with simultaneous locomotion, pupil area, facial movement (FaceMap PC1) and an example raw ECoG trace shown below. **b**, Average spatial maps (n = 6) showing parcel-wise differences in ACh3.0 and jRCaMP1b activity upon transition to high facial motion (top) and locomotion (bottom), showing peak ΔF/F values at 0 to 1s post high facial motion or locomotion onset. **c**, Peak (at 0 to 1s from state transition) ΔF/F difference values (post – pre onset) from parcels in the left hemisphere are plotted against their anterior-to-posterior position based on center of mass. r_s_ indicates Spearman’s rank order correlation coefficient for correlation between mean value and anterior-posterior rank across parcels. Line indicates linear fit for visualization. **d**, Same as in (c) for ΔF/F difference values (mean +/- SEM) for sustained states (high-low facial motion PC1 onset, top and locomotion-high facial motion, bottom).

**Figure S6.**
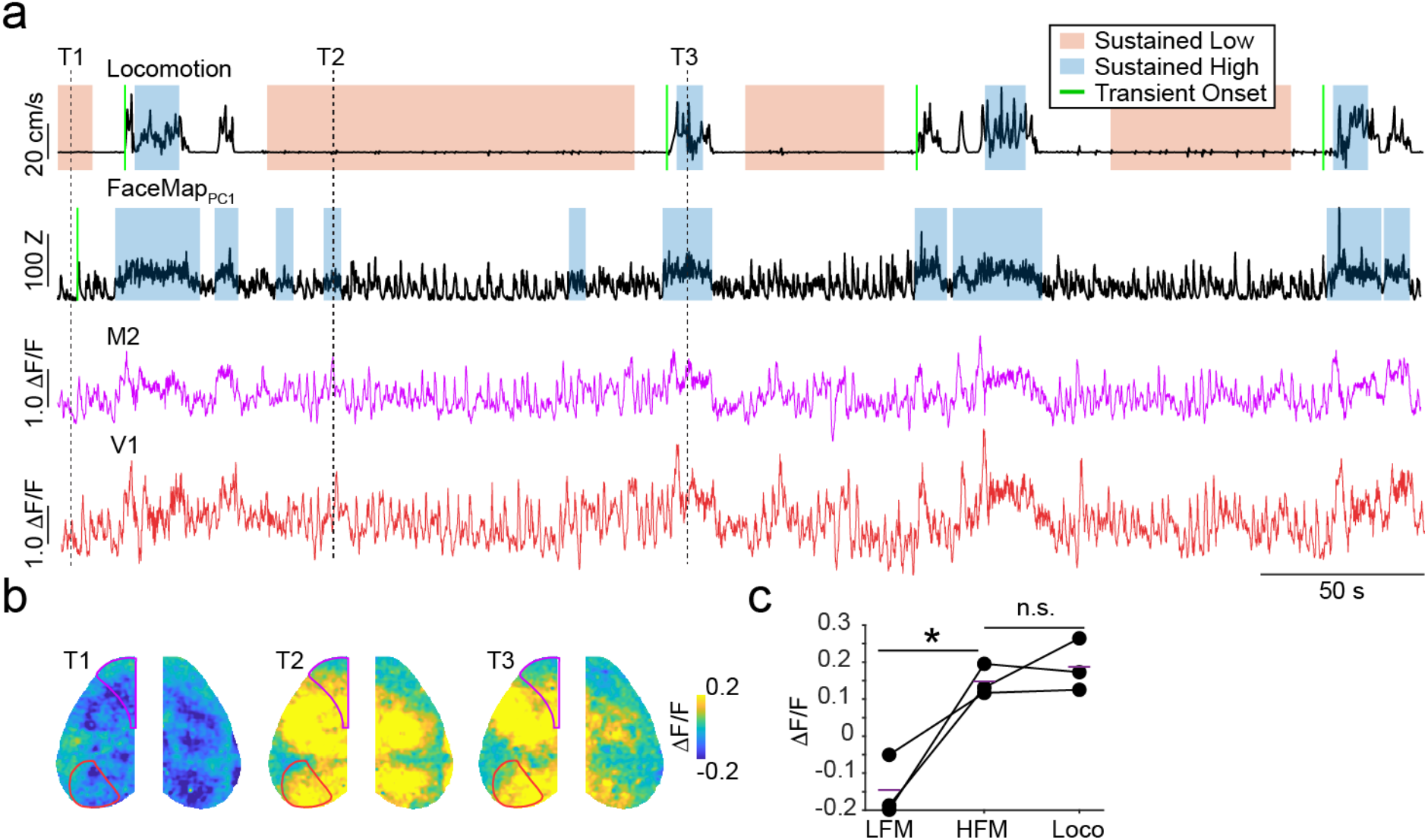
Cholinergic signal in ChAT-GCaMP mice during movement-defined behavioral states. **a**, Example time series showing behavioral measures (locomotion and FaceMap PC1) and GCaMP6 signals from the M2 (purple) and V1 (red) parcels in a Chat-Cre^+/0^Ai162^F/0^ mouse. **b**, Representative image frames from a period of no locomotion and low facial motion activity (T1), no locomotion and high facial motion activity (T1), and locomotion and high facial motion activity (T3). **c**, Individual animal values and population mean whole cortex ΔF/F values during sustained low facial motion, high facial motion, and locomotion states. Inset shows individual and population mean ΔF/F difference values between high-low facial motion and locomotion-high facial motion. *indicates p < 0.05 for post-hoc LSD comparisons following repeated measures ANOVA.

## Materials and Methods

### Animals

Male and female C57BL/6J, ChAT-Cre (B6J.ChAT-IRES-Cre::frt-neo-frt, Jax stock no. 028861) and Ai162 (TIT2L-GCaMP6s-ICL-tTA2, Jax stock no. 031562) mice were kept on a 12h light/dark cycle, provided with food and water ad libitum, and housed individually following headpost implants. Imaging experiments were performed during the light phase of the cycle. All animal handling and experiments were performed according to the ethical guidelines of the Institutional Animal Care and Use Committee of the Yale University School of Medicine.

### Neonatal sinus injections

Brain-wide expression of the ACh sensor ACh3.0 and the calcium indicator jRCaMP1b was achieved via postnatal sinus injection^11, 39^. Specifically, P0-P1 litters were removed from their home cage and placed on a heating pad. Pups were kept on ice for 5 min to induce anesthesia via hypothermia and then maintained on a metal plate surrounded by ice for the duration of the injection. Under a dissecting microscope, two small incisions were made in the skin over the transverse sinuses. Viral injections were made with a NanoFil (WPI) attached to a 36-gauge needle and an UltraMicroPump (WPI) mounted to a stereotaxic arm. The needle was slowly lowered through the skull into the underlying transverse sinus. Pups were injected bilaterally with 2 μl of AAV9-hsyn-ACh3.0 (1.8×10^13 gc/ml) and 2 μl of AAV9-hsyn-NES-jRCaMP1b (2.5×10^13 gc/ml, Addgene) per hemisphere. The hSyn promoter drives expression in most excitatory and inhibitory neurons. Subset of pups were bilaterally injected with 4 μl AAV9-hsyn-EGFP (3.4×10^13 gc/ml, Addgene) or 4 μl ACh3.0 (1.8×10^13 gc/ml) per hemisphere. Viruses were injected at 10 nl/s, and the needle was left in the sinus for 30 s following the injection. Incision sites were sealed with Vetbond glue, and pups were moved to a heating pad. Once the entire litter was injected, pups were gently rubbed with home cage bedding and nesting material and returned to their home cage. Chat-Cre^+/0^Ai162^F/0^ ^67^ mice endogenously expressed GCamp6s in cholinergic neurons and terminals and were not injected with virus.

### Surgical procedures

All surgical implant procedures were performed on adult mice (>P50). Mice were anesthetized using 1-2% isoflurane and maintained at 37°C for the duration of the surgery. The skin and fascia above the skull were removed from the nasal bone to the posterior of the intraparietal bone and laterally between the temporal muscles. The surface of the skull was thoroughly cleaned with saline and the edges of the incision secured to the skull with Vetbond. A custom titanium headpost was secured to the posterior of the nasal bone with transparent dental cement (Metabond, Parkell), and a thin layer of dental cement was applied to the entire dorsal surface of the skull. Next, a layer of cyanoacrylate (Maxi-Cure, Bob Smith Industries) was used to cover the skull and left to cure ∼30 min at room temperature to provide a smooth surface for transcranial imaging.

For simultaneous ECoG implants, a dental drill was used to make a small craniotomy (∼1mm) lateral to V1 in the right hemisphere. A silver ball electrode was placed in the craniotomy and a teflon coated silver reference wire implanted in the contralateral cerebellar hemisphere. A ground wire was wrapped around a skull screw placed in the ipsilateral intraparietal bone. For simultaneous basal forebrain stimulation in a subset of animals, stainless steel bipolar stimulating electrodes (125 um diameter, Invivo1) were implanted at the following coordinates (AP = -0.5 mm, ML = 1.6 mm, DV = 4 mm, angle = 0 degrees or AP = -0.5 mm, ML = 3.3 mm, DV = 3.7 mm, angle = 20 degrees) to target the nucleus basalis in the right hemisphere. Stimulation comprised a brief burst at 100 Hz (1 ms pulse width, 20 pulses, 60-100 uA). Desynchronization in cortical ECoG was used to verify correct electrode placement and to titrate the stimulation intensity.

### Widefield imaging

Widefield calcium and cholinergic imaging was performed using a Zeiss Axiozoom with a PlanNeoFluar Z 1x, 0.25 numerical aperture objective with a 56 mm working distance. Epifluorescent excitation was provided by an LED bank (Spectra X Light Engine, Lumencor) using three output wavelengths: 395/25, 470/24, and 575/25 nm. Emitted light passed through a dual camera image splitter (TwinCam, Cairn Research) then through either a 525/50 or 630/75 emission filter (Chroma) before it reached two sCMOS cameras (Orca-Flash V3, Hamamatsu). Images were acquired at 512×512 resolution after 4x pixel binning, and each channel was acquired at 10 Hz with 20 ms exposure. Images were saved to a solid-state drive using HCImage software (Hamamatsu).

Backscatter illumination was provided by LEDs (Thorlabs M530L4, M625L4) centered and narrowly filtered at 530 nm and 625 nm coupled to a 1000 um diameter bifurcated fiber (BFY1000LS02) that terminated 45 degrees incident to the brain surface. Image frames capturing backscatter at 530 and 625 nm were acquired at 10 Hz and interleaved with the usual fluorescence emission acquisition.

All imaging was performed during the second half of the light cycle in awake, behaving mice that were head-fixed so that they could freely run on a cylindrical wheel^3, 4, 68^. A magnetic angle sensor (Digikey) attached to the wheel continuously monitored wheel motion. Mice received at least three wheel-training habituation sessions before imaging to ensure consistent running bouts (see Figure S4). During widefield imaging sessions, the face (including the pupil and whiskers) was illuminated with an IR LED bank and imaged with a miniature CMOS camera (Blackfly s-USB3, Flir) with a frame rate of 10 Hz. To monitor the ECoG, we used a DP-311A differential amplifier with active headstage (Warner Instruments). Signals were amplified 1000x, filtered between 0.1Hz and 1000 Hz, and digitized at 5000 Hz using a Power 1401 acquisition board (CED).

### Visual stimulation and air-puffs

Small (40 degree diameter for backscatter sessions and 20 degree diameter for other sessions) sinusoidal drifting gratings (2 Hz, 0.04 cycles/degree, 100% contrast) were generated using Psychtoolbox in Matlab and presented on an LCD monitor at a distance of 20 cm from the right eye. Stimuli were presented for 2 seconds with a 5 second inter-stimulus interval. Air-puff stimuli were delivered using a thin metal tube aimed at the fur along the back and coupled to a solenoid valve (Clark Solutions) that delivered brief (200 ms) puffs of compressed air.

### Histology

Histological validation was performed on a subset of animals at the conclusion of imaging experiments. Mice were deeply anesthetized with isoflurane and perfused transcardially with phosphate buffered saline (PBS) followed by 4 percent paraformaldehyde in PBS. Brains were postfixed overnight at 4 degrees and embedded in 1% agarose, and 50 um sagittal sections were cut on a vibratome (VT1000, Leica). Slices were pretreated in blocking solution (2 percent normal goat serum, 0.1 percent Triton X-100 in PBS) for four hours then incubated with primary antibodies at 1:1000 (rabbit anti-GFP and guinea pig anti-NeuN, Invitrogen) at 4 degrees for 24 hours. The following day, slices were washed with PBS and incubated in secondary antibodies (anti-rabbit-Alexa Fluor 488 and anti-guinea pig-Alexa Fluor 647, Invitrogen) at 1:1000 for two hours at room temperature. Slices were washed 3 times in PBS then mounted on glass slides in Vectashield antifade mounting medium (Vector Laboratories). Epifluorescent images were acquired on an Olympus BX53 fluorescence microscope. Confocal images were taken with a Zeiss LSM 710.

### *ACh3.0* imaging *ex vivo*

Under isoflurane anesthesia, mice were decapitated and coronal slices (∼300 um thick) were cut in ice-cold external solution containing (in mM): 100 choline chloride, 25 NaHCO3, 1.25 NaH2PO4, 2.5 KCl, 7 MgCl2,

0.5 CaCl2, 15 glucose, 11.6 sodium ascorbate and 3.1 sodium pyruvate, bubbled with 95% O2 and 5% CO2. Slices were transferred to artificial cerebrospinal fluid (ACSF) containing (in mM): 127 NaCl, 25 NaHCO3, 1.25 NaH2PO4, 2.5 KCl, 1 MgCl2, 2 CaCl2 and 15 glucose, bubbled with 95% O2 and 5% CO2. After an incubation period, slices were moved to a modified recording chamber under the objective of the widefield microscope and constantly perfused with oxygenated ACSF. Slices were imaged using the same protocol as during *in vivo* imaging sessions (10 Hz, 20 ms exposure). A glass pipette filled with 20 millimolar carbachol was mounted in a micromanipulator and lowered to just above the slice. A Picospritzer (Parker Hannifin Corp) was used to deliver 200 ms puffs of carbachol. During some trials, scopolamine (20 uM) was added to the ACSF before carbachol puffs were applied.

### Data analysis

All analyses were conducted using custom-written scripts in MATLAB (Mathworks) and Prism9 (GraphPad). All statistical results are listed in Supplemental Table 2. Exact correlation values and p-values for Figures 4 and 5 are shown in Supplemental Table 3.

#### Preprocessing of imaging data

512×512 images were down-sampled to 256×256, and frames were grouped by excitation wavelength (395 nm, 470 nm, 575 nm). For dual color imaging, green and red images were acquired using different cameras and registered via automatic ‘rigid’ transformation using imregtform in MATLAB. In some cases, registration points were manually selected and a ‘similarity’ geometric transformation was applied. Images were then detrended and baseline corrected to calculate ΔF/F images. Specifically, for each pixel, a 100th order fir1 filter with 0.001 Hz frequency cutoff was applied to extract the low pass filtered signal. This low pass signal was used as baseline (F0), and ΔF/F for each pixel was calculated as (F-F0)/F0, where F is the raw unfiltered signal. ΔF/F values were used for all subsequent analyses. Images were registered to the Allen Common Coordinate framework (CCFv3) using manually selected control points and ‘similarity’ based geometric transformation. Time series for individual brain parcels were then extracted by averaging ΔF/F values across all pixels within a parcel.

#### Hemodynamic correction

Common methods for correction of hemodynamic artifacts are typically based on linear regression of a neural activity-dependent signal against an activity-independent signal and performed on a pixel-by-pixel basis, ignoring the spatial correlations among neighboring pixels that exist within and between the two signals. However, accounting for such correlations can be advantageous for mitigating the effects of noise. We experimentally confirmed that excitation of ACh3.0 with 395 nm light results in fluorescence with substantially reduced dependence on ACh (Figure S2), enabling us to use this approach to correct the signal collected with 470 nm excitation. We now present our mathematical formulation for hemodynamic artifact removal as the optimal linear predictor for the neuronal time series, given the 470 nm- and 395 nm-excitation signals.

Let *y*_1_ and *y*_2_ be *p* × 1 random signals corresponding to *p* pixels of the 470 nm and 395 nm signals, respectively. Let *x* and *z* be mutually uncorrelated *p* × 1 random signals corresponding to *p* pixels of the ACh-dependent and -independent (hemodynamic) signals, respectively. We consider the following linear model:

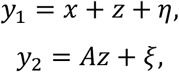

where *η* and *ξ* are white Gaussian *p* × 1 noise signals and A is an unknown *p* × *p* real invertible matrix. Given the above-mentioned model, our goal is to estimate the signal *x*.

It can be verified that the optimal linear estimator of *x* in the sense of Minimum Mean Squared Error (MMSE) is given by:

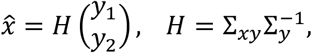

Where 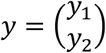 is given by stacking *y*_*1*_ on top of *y*_*2*_, Σ_*y*_ = 𝔼[*yy*^*T*^] is the autocorrelation matrix of *y*, and Σ_*xy*_ = 𝔼[*xy*^*T*^] is the cross-correlation matrix between *x* and *y*. While Σ_*y*_ can be estimated directly from the observations, this is not trivially the case for Σ_*xy*_ since it depends on the unknown signal *x*. Nevertheless, we show below that Σ_*xy*_ can be expressed as:

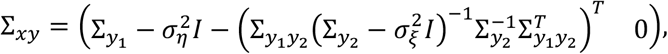

Where 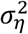 and 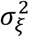 are the noise variances of *η* and *ξ*, respectively, and *I* is the *p* × *p* identity matrix. Importantly, all quantities in the formula for Σ_*xy*_ can be estimated directly from the observations of *y*_1_ and *y*_2_. The noise variances 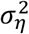 and 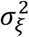 were evaluated according to the median of the singular values of the corresponding correlation matrices 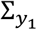 and 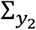.

Proof for the formula for Σ_*xy*_:

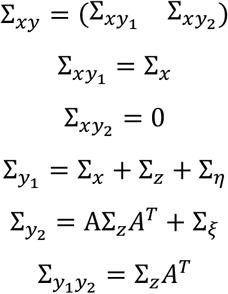

Using these equations, we can extract 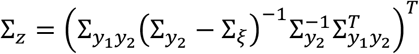 which leads to:

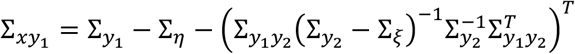

Note that in the case of a single pixel, i.e. *p* = 1, our estimator reduces to a simple regression based on pixelwise correlations. However, taking *p* to be all pixels at once is both computationally expensive and unnecessary. Therefore, we performed these evaluations locally, where the time trace of each pixel was computed based on a patch of its nearest spatial neighbors. The size of the patch was determined according to the amount of time samples available. Using a bigger patch would lead to bigger covariance matrices to be estimated and therefore requires longer time traces. For our sessions a patch size of 9 × 9 was empirically selected as most adequate.

#### Validation of hemodynamics correction

The novel spatial regression approach used here was validated by comparing results with other known hemodynamics correction strategies. Several benchmarks were used to determine the effectiveness of our method. First, we and others have observed that strong stimuli, such as a high-contrast visual stimulus, evoke negative dips in fluorescence signals that are driven by hemodynamic photon absorption ^41, 42^. Here, we compared this visual stimulus-evoked negativity in V1 for uncorrected ΔF/F signals (measured for 470 nm excitation) and ΔF/F signals corrected by either spatial or pixel-wise regression of signals collected at 395 nm excitation. Visual trials were normalized by subtracting the baseline mean in the -2 to 0 s pre-stimulus period, and the minimum normalized ΔF/F values at 0 to 5 s after stimulus onset were calculated for each trial. These values were then averaged across all trials in an animal, and statistical comparisons were made within mice between the three methods using protected paired t-tests following repeated measures ANOVA. A complementary approach involves the measurement of hemodynamic absorption from changes in the reflectance (“backscatter”) of green photons while simultaneously performing fluorescence imaging ^41, 42^. Here, we carried out pixel-wise and spatial regression of Ach 3.0 signals using 530 nm backscatter data and compared results with 395 nm-based correction. Finally, we also assessed the efficiency of the different correction approaches to reduce the variance in ΔF/F signals collected from GFP-expressing control mice, where all fluctuations are assumed to result primarily from hemodynamic variation. Statistical comparisons on ΔF/F maximum negativity values post-air-puff were performed in a similar manner as descried above for visual stimuli.

#### Preprocessing of behavior data

Facial movements were extracted from face videography using FaceMap^9^. The toolbox applies singular value decomposition (SVD) to the face movie to extract the 1000 dimensions or principal components (PCs) that explain the distinct movements apparent on the mouse’s face. Here, we included the top 25 PCs for regression analyses and the first PC for other analyses. To extract face movement states, we focused on quiescence periods only, because locomotion is consistently associated with increased facial motion. First, PC1 data were Z-score normalized within a session, and the high/low thresholds corresponding to 60% and 40% quantiles in the data distribution during quiescence only were extracted for that session. Data were then smoothed using a 1s-window moving average filter, and epochs during which smoothed data continuously exceeded the high/low Z-threshold for at least 0.5 s were considered high/low face state, respectively.

Wheel position was determined from the output of the linear angle detector. The circular wheel position variable was first transformed to the [-π,π] interval. The phases were then circularly unwrapped to get running distance as a linear variable, and locomotion speed was computed as a differential of distance (cm/s). A change-point detection algorithm detected locomotion onset/offset times based on changes in standard deviation of speed. First, moving standard deviations of speed were computed in 2 s windows, which was the temporal resolution of the changepoint analysis. Initial locomotion onset/offset times were then estimated from periods when the moving standard deviations exceeded/ fell below an empirical threshold of 0.005. These estimates were refined by computing the time points corresponding to the maximum of the moving forward/backward Z-score in 2 s windows around each onset/offset time. Locomotion epochs that were too short (<1s) or where average speed was too low (<2 cm/s) were removed.

### Post-processing analysis

Imaging data from individual parcels were used for subsequent analyses. Some medial Allen parcels (PL, ACA, RSPv) were masked out because of prominent vasculature along the midline. Colliculus regions (SC, IC) were also removed, resulting in final 23 Allen CCFv3 parcels per hemisphere (see Figure S2). The primary visual cortex (V1 or VlSp) and the secondary motor area in the frontal cortex (M2 or Mos) were selected as representative areas for some population summaries and statistical comparisons. The anterior-posterior spatial ranking of Allen CCFv3 areas was determined by calculating the relative centers of mass for each parcel (see Figure S2). The ordering was as follows (from anterior to posterior): Mos, Mop, SSp-m, SSp-ul, SSp-n, SSp-ll, SSp-un, SSp-tr, SSp-bfd, SSs, PTLp, VlSam, AUD, RSPagl, RSPd, VlSal, VlSpm, Tea, VlSli, VlSl, VlSp, VlSpor, VlSpl.

#### Basal Forebrain Stimulation-Evoked Activity

Trials where basal forebrain stimulation caused locomotion were automatically excluded. Data were normalized by subtracting the baseline mean in the pre-stimulus period -2 to 0 s. Difference ΔF/F data were averaged across all trials within an animal and summarized as mean +/- standard error of mean across animals.

#### Regression with behavioral variables

Linear regression of continuous data from full imaging sessions was used to predict ACh3.0 and jRCaMP1b activity in each parcel using behavioral variables as predictors. 10 fold cross validation was performed by partitioning the time series into 10 chunks and using independent test partitions in each fold. The prediction accuracy was calculated on testing data in each of the 10 folds and averaged to get a final cross-validated R^2^. Individual regression models were built using 25 FaceMap PCs, pupil area, or wheel speed. A two-way repeated measures ANOVA was used to assess whether cross validated R^2^ values are significantly different across brain areas and behavioral states, followed by post-hoc LSD test for states. Spearman’s rank-order correlation was used to assess the relationship between mean cross validated R^2^ values and Allen-defined parcels’ anterior-posterior positioning.

#### State Transitions and Sustained States

Behavioral states were categorized into high/low states (high/low face or locomotion/quiescence) based on movement as described above (see preprocessing of Behavior Data). Changes at transition from low to high movement state (face onset or locomotion onset) were defined as state transients (see Figure 3a). For locomotion onset, only locomotion trials that contained at least 5 s of running and were preceded by a minimum 10 s of quiescence were included. For high face state onset, if a high face state was preceded by at least 4 s of a low face state, the transition between the two states was used as the high state onset time point. For each animal, state transients were quantified by first subtracting mean pre-onset (−5 to -3s for locomotion and -4 to - 2s for face) from post-onset ΔF/F values and averaging data across all state transitions, then calculating the peak ΔF/F value at 0 to 1 s after onset. Whole cortex-averaged transients were obtained by averaging peak ΔF/F values across all parcels in one hemisphere, and a one-sample Student’s t-test was conducted to test significant differences from zero. A two-way repeated measures ANOVA was used to assess any significant differences in state transients across brain areas and behavioral states. Spearman’s rank-order correlation was used to assess the relationship between mean peak ΔF/F values and Allen-defined parcels’ anterior-posterior positioning.

Sustained behavioral states were defined as high or low movement states that persisted for at least 5 s (see Figure 3a). For sustained locomotion states, it was required that locomotion started at least 3 s before and ended at least 3 s after the state boundaries. Similarly, for sustained quiescence states, it was required that any locomotion did not occur at least 10 s before and 10 s after a given period. Sustained high/low face states were extracted during quiescence only and were required to be a minimum of 5 s in duration. State data were thus compared between low FaceMap, high FaceMap, and locomotion states. To ensure accurate comparison between states, the number of state epochs and total duration within each epoch were matched across states within each session. ACh3.0, RCaMP1b, and Chat-Gcamp6s ΔF/F responses during these three states were quantified for each parcel, and whole cortex-averaged responses were obtained by averaging ΔF/F values across all parcels in one hemisphere. A two-way repeated measures ANOVA was used to assess whether ΔF/F responses varied across brain areas and behavioral states, followed by post-hoc LSD tests to test differences between high and low FaceMap states as well as locomotion and high FaceMap state. The response to facial movement and locomotion were quantified as difference between high and low FaceMap ΔF/F values and between locomotion and high FaceMap ΔF/F values, respectively. Spearman’s rank-order correlation was used to assess the relationship between difference ΔF/F values and anterior-posterior rank of Allen-defined parcels.

#### ECoG processing

The multitaper spectrogram in Figure 1 was estimated from the raw ECOG signal using the Chronux toolbox (using sliding windows of 3 s with 0.5 s overlap). ECoG power was calculated for each behavioral state by estimating the bandpower in high (30-100 Hz) and low frequency (1-10 Hz) bands. The ECoG high and low frequency power values were compared between high versus low FaceMap and locomotion versus high FaceMap states using Student’s paired t-tests.

#### State-dependent correlations

Time-lagged correlations between every pair of Allen CCF parcels in Ach 3.0 or jRCaMP1b signals were computed by performing cross-correlations with 500 ms time lag and extracting the maximum correlation coefficient. The resulting parcel-wise correlation matrices were calculated during sustained behavioral states with a minimum duration of 5 s to allow enough time frames for lagged correlations. To ensure accurate comparison in correlations between states, the number of state epochs and total duration within each epoch were matched for locomotion, high, and low facial movement states within each session. Correlation matrices were calculated for each state epoch and averaged all epochs and sessions for each animal to get one correlation matrix per animal per state. Cross-correlations between Ach 3.0 and jRCaMP1b activity for distinct behavioral states were calculated in a similar manner except the correlations were performed between the two signals within a parcel. Correlation matrices were compared between locomotion versus high facial motion states as well as high facial motion versus low facial motion states. To assess the significance of differences in pairwise correlations between states (high versus low facial motion and locomotion versus high facial motion), a stratified permutation test was used. In each permutation, state labels were shuffled across epochs within each animal, and the corresponding correlation matrices were averaged across all epochs of a particular state to get one permuted correlation matrix per state per animal. These values were then averaged across animals to get a mean difference between states. This was repeated 10,000 times to build a null distribution of mean differences which was compared against the observed mean difference to determine p-values. Multiple comparisons correction was performed by setting the false discovery rate (FDR) to q < 0.01 and using Benjamini-Yekutieli method in fdr_bh.m toolbox in MATLAB. Correlation values and adjusted p-values are available in the Supplemental Table 3.

---

**Supplemental Table 1:**
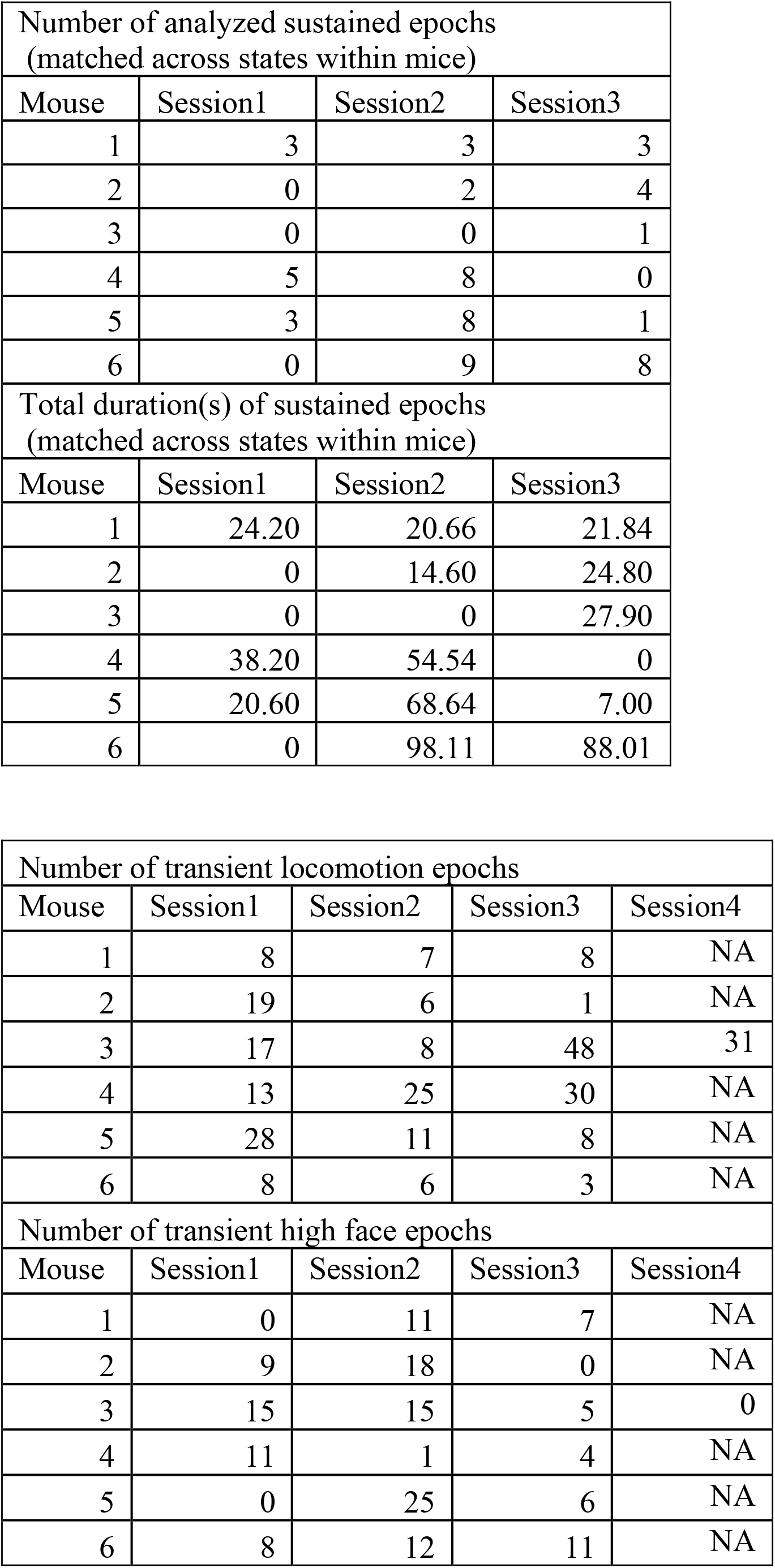
State epoch number and duration

**Supplemental Table 2.**
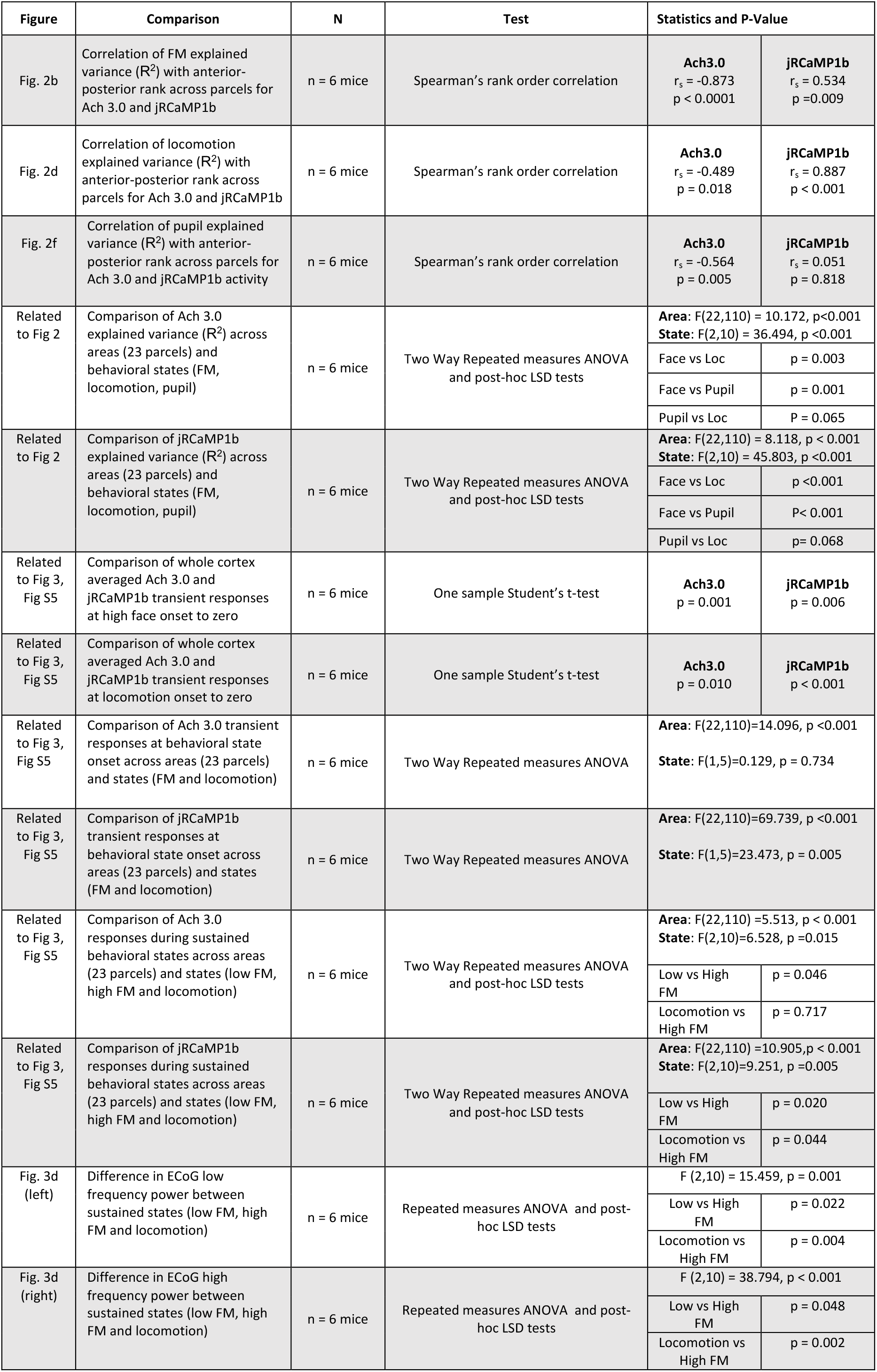
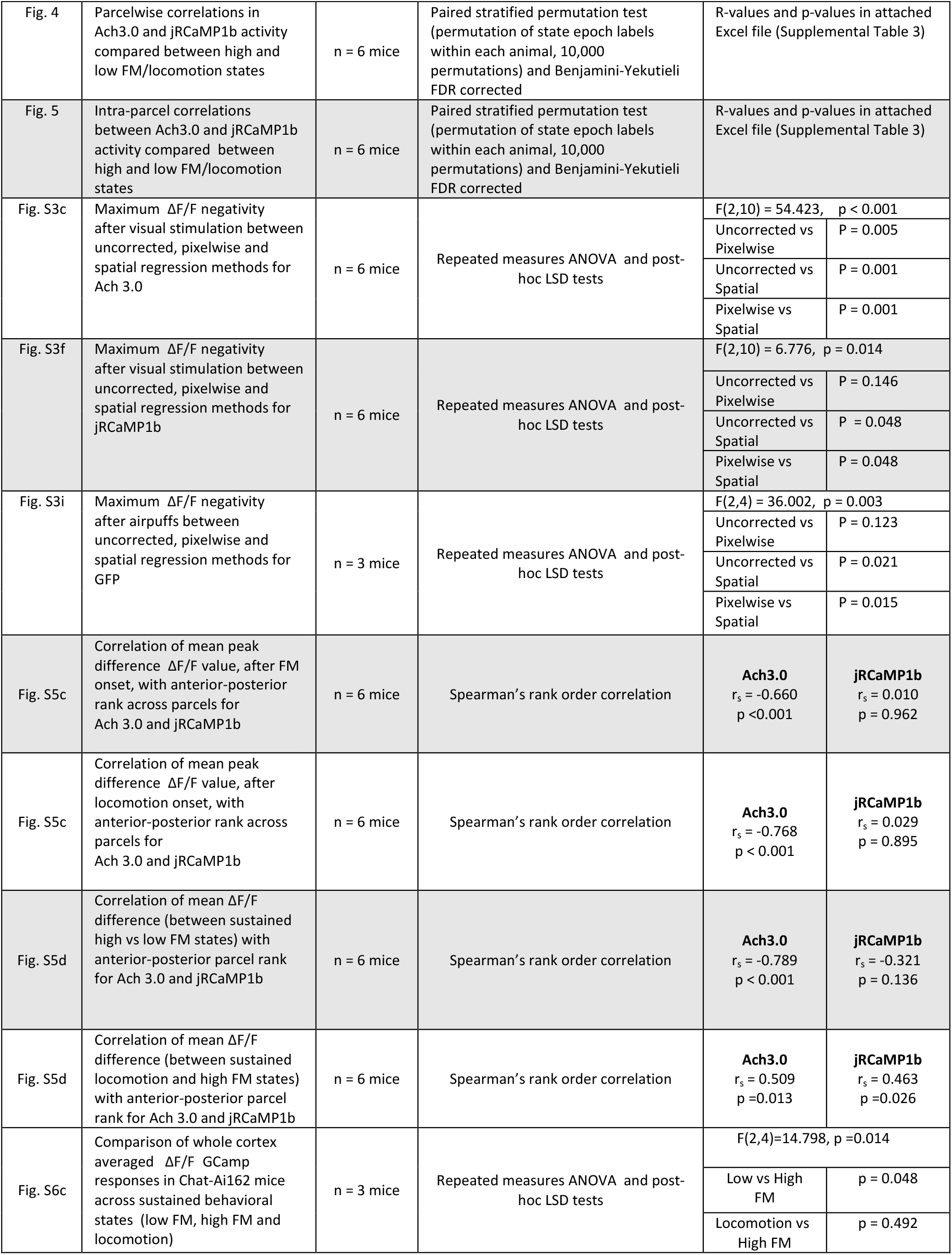
Summary of all statistical analyses.

| Figure | Comparison | N | Test | Statistics and P-Value |  |
| --- | --- | --- | --- | --- | --- |
| Fig. 2b | Correlation of FM explained variance ($R^2$) with anterior-posterior rank across parcels for Ach 3.0 and jRCaMP1b | n = 6 mice | Spearman's rank order correlation | **Ach3.0**<br>$r_s = -0.873$<br>$p < 0.0001$ | **jRCaMP1b**<br>$r_s = 0.534$<br>$p = 0.009$ |
| Fig. 2d | Correlation of locomotion explained variance ($R^2$) with anterior-posterior rank across parcels for Ach 3.0 and jRCaMP1b | n = 6 mice | Spearman's rank order correlation | **Ach3.0**<br>$r_s = -0.489$<br>$p = 0.018$ | **jRCaMP1b**<br>$r_s = 0.887$<br>$p < 0.001$ |
| Fig. 2f | Correlation of pupil explained variance ($R^2$) with anterior-posterior rank across parcels for Ach 3.0 and jRCaMP1b activity | n = 6 mice | Spearman's rank order correlation | **Ach3.0**<br>$r_s = -0.564$<br>$p = 0.005$ | **jRCaMP1b**<br>$r_s = 0.051$<br>$p = 0.818$ |
| Related to Fig 2 | Comparison of Ach 3.0 explained variance ($R^2$) across areas (23 parcels) and behavioral states (FM, locomotion, pupil) | n = 6 mice | Two Way Repeated measures ANOVA and post-hoc LSD tests | **Area:** F(22,110) = 10.172, $p < 0.001$<br>**State:** F(2,10) = 36.494, $p < 0.001$ | |
| | | | | Face vs Loc | $p = 0.003$ |
| | | | | Face vs Pupil | $p = 0.001$ |
| | | | | Pupil vs Loc | $P = 0.065$ |
| Related to Fig 2 | Comparison of jRCaMP1b explained variance ($R^2$) across areas (23 parcels) and behavioral states (FM, locomotion, pupil) | n = 6 mice | Two Way Repeated measures ANOVA and post-hoc LSD tests | **Area:** F(22,110) = 8.118, $p < 0.001$<br>**State:** F(2,10) = 45.803, $p < 0.001$ | |
| | | | | Face vs Loc | $p < 0.001$ |
| | | | | Face vs Pupil | $P < 0.001$ |
| | | | | Pupil vs Loc | $p = 0.068$ |
| Related to Fig 3, Fig S5 | Comparison of whole cortex averaged Ach 3.0 and jRCaMP1b transient responses at high face onset to zero | n = 6 mice | One sample Student's t-test | **Ach3.0**<br>$p = 0.001$ | **jRCaMP1b**<br>$p = 0.006$ |
| Related to Fig 3, Fig S5 | Comparison of whole cortex averaged Ach 3.0 and jRCaMP1b transient responses at locomotion onset to zero | n = 6 mice | One sample Student's t-test | **Ach3.0**<br>$p = 0.010$ | **jRCaMP1b**<br>$p < 0.001$ |
| Related to Fig 3, Fig S5 | Comparison of Ach 3.0 transient responses at behavioral state onset across areas (23 parcels) and states (FM and locomotion) | n = 6 mice | Two Way Repeated measures ANOVA | **Area:** F(22,110)=14.096, $p < 0.001$<br>**State:** F(1,5)=0.129, $p = 0.734$ | |
| Related to Fig 3, Fig S5 | Comparison of jRCaMP1b transient responses at behavioral state onset across areas (23 parcels) and states (FM and locomotion) | n = 6 mice | Two Way Repeated measures ANOVA | **Area:** F(22,110)=69.739, $p < 0.001$<br>**State:** F(1,5)=23.473, $p = 0.005$ | |
| Related to Fig 3, Fig S5 | Comparison of Ach 3.0 responses during sustained behavioral states across areas (23 parcels) and states (low FM, high FM and locomotion) | n = 6 mice | Two Way Repeated measures ANOVA and post-hoc LSD tests | **Area:** F(22,110) = 5.513, $p < 0.001$<br>**State:** F(2,10)=6.528, $p = 0.015$ | |
| | | | | Low vs High FM | $p = 0.046$ |
| | | | | Locomotion vs High FM | $p = 0.717$ |
| Related to Fig 3, Fig S5 | Comparison of jRCaMP1b responses during sustained behavioral states across areas (23 parcels) and states (low FM, high FM and locomotion) | n = 6 mice | Two Way Repeated measures ANOVA and post-hoc LSD tests | **Area:** F(22,110) = 10.905, $p < 0.001$<br>**State:** F(2,10)=9.251, $p = 0.005$ | |
| | | | | Low vs High FM | $p = 0.020$ |
| | | | | Locomotion vs High FM | $p = 0.044$ |
| Fig. 3d (left) | Difference in ECoG low frequency power between sustained states (low FM, high FM and locomotion) | n = 6 mice | Repeated measures ANOVA and post-hoc LSD tests | F (2,10) = 15.459, $p = 0.001$ | |
| | | | | Low vs High FM | $p = 0.022$ |
| | | | | Locomotion vs High FM | $p = 0.004$ |
| Fig. 3d (right) | Difference in ECoG high frequency power between sustained states (low FM, high FM and locomotion) | n = 6 mice | Repeated measures ANOVA and post-hoc LSD tests | F (2,10) = 38.794, $p < 0.001$ | |
| | | | | Low vs High FM | $p = 0.048$ |
| | | | | Locomotion vs High FM | $p = 0.002$ |
| Fig. 4 | Parcelwise correlations in Ach3.0 and jRCaMP1b activity compared between high and low FM/locomotion states | n = 6 mice | Paired stratified permutation test (permutation of state epoch labels within each animal, 10,000 permutations) and Benjamini-Yekutieli FDR corrected | R-values and p-values in attached Excel file (Supplemental Table 3) |  |
| Fig. 5 | Intra-parcel correlations between Ach3.0 and jRCaMP1b activity compared between high and low FM/locomotion states | n = 6 mice | Paired stratified permutation test (permutation of state epoch labels within each animal, 10,000 permutations) and Benjamini-Yekutieli FDR corrected | R-values and p-values in attached Excel file (Supplemental Table 3) |  |
| Fig. S3c | Maximum $\Delta F/F$ negativity after visual stimulation between uncorrected, pixelwise and spatial regression methods for Ach 3.0 | n = 6 mice | Repeated measures ANOVA and post-hoc LSD tests | F(2,10) = 54.423, p < 0.001 | |
|  |  |  |  | Uncorrected vs Pixelwise | P = 0.005 |
|  |  |  |  | Uncorrected vs Spatial | P = 0.001 |
|  |  |  |  | Pixelwise vs Spatial | P = 0.001 |
| Fig. S3f | Maximum $\Delta F/F$ negativity after visual stimulation between uncorrected, pixelwise and spatial regression methods for jRCaMP1b | n = 6 mice | Repeated measures ANOVA and post-hoc LSD tests | F(2,10) = 6.776, p = 0.014 | |
|  |  |  |  | Uncorrected vs Pixelwise | P = 0.146 |
|  |  |  |  | Uncorrected vs Spatial | P = 0.048 |
|  |  |  |  | Pixelwise vs Spatial | P = 0.048 |
| Fig. S3i | Maximum $\Delta F/F$ negativity after airpuffs between uncorrected, pixelwise and spatial regression methods for GFP | n = 3 mice | Repeated measures ANOVA and post-hoc LSD tests | F(2,4) = 36.002, p = 0.003 | |
|  |  |  |  | Uncorrected vs Pixelwise | P = 0.123 |
|  |  |  |  | Uncorrected vs Spatial | P = 0.021 |
|  |  |  |  | Pixelwise vs Spatial | P = 0.015 |
| Fig. S5c | Correlation of mean peak difference $\Delta F/F$ value, after FM onset, with anterior-posterior rank across parcels for Ach 3.0 and jRCaMP1b | n = 6 mice | Spearman's rank order correlation | **Ach3.0**<br>$r_s = -0.660$<br>p < 0.001 | **jRCaMP1b**<br>$r_s = 0.010$<br>p = 0.962 |
| Fig. S5c | Correlation of mean peak difference $\Delta F/F$ value, after locomotion onset, with anterior-posterior rank across parcels for Ach 3.0 and jRCaMP1b | n = 6 mice | Spearman's rank order correlation | **Ach3.0**<br>$r_s = -0.768$<br>p < 0.001 | **jRCaMP1b**<br>$r_s = 0.029$<br>p = 0.895 |
| Fig. S5d | Correlation of mean $\Delta F/F$ difference (between sustained high vs low FM states) with anterior-posterior parcel rank for Ach 3.0 and jRCaMP1b | n = 6 mice | Spearman's rank order correlation | **Ach3.0**<br>$r_s = -0.789$<br>p < 0.001 | **jRCaMP1b**<br>$r_s = -0.321$<br>p = 0.136 |
| Fig. S5d | Correlation of mean $\Delta F/F$ difference (between sustained locomotion and high FM states) with anterior-posterior parcel rank for Ach 3.0 and jRCaMP1b | n = 6 mice | Spearman's rank order correlation | **Ach3.0**<br>$r_s = 0.509$<br>p = 0.013 | **jRCaMP1b**<br>$r_s = 0.463$<br>p = 0.026 |
| Fig. S6c | Comparison of whole cortex averaged $\Delta F/F$ GCaMP responses in Chat-Ai162 mice across sustained behavioral states (low FM, high FM and locomotion) | n = 3 mice | Repeated measures ANOVA and post-hoc LSD tests | F(2,4)=14.798, p=0.014 | |
|  |  |  |  | Low vs High FM | p = 0.048 |
|  |  |  |  | Locomotion vs High FM | p = 0.492 |

